# Secreted spermidine synthase reveals a paracrine role for PGC1α-induced growth suppression in prostate cancer

**DOI:** 10.1101/2024.10.04.614869

**Authors:** Ariane Schaub-Clerigué, Ivana Hermanova, Ainara Pintor-Rial, Alice Macchia, Lorea Valcarcel-Jimenez, Benoit Lectez, Saioa Garcia-Longarte, Maider Fagoaga-Eugui, Amaia Zabala-Letona, Mikel Pujana-Vaquerizo, Félix Royo, Mikel Azkargorta, Edurne Berra, James D. Sutherland, Héctor Peinado, Juan Manuel Falcón-Perez, Félix Elortza, Arkaitz Carracedo, Verónica Torrano

## Abstract

Prostate cancer is the fifth cause of death by cancer worldwide, second in incidence in the male population. The definition of the molecular basis of its development and the oncogenic signals driving lethality continue to be important objectives in prostate cancer research. Prior work from others and us has demonstrated that loss of PGC1α expression results in a metabolic, signaling and transcriptional reprogramming that supports the development of metastatic disease. However, we do not fully understand the spectrum of tumor suppressive effects regulated by this co-regulator. Here we show that PGC1α governs non-cell autonomous paracrine tumor suppression in prostate cancer. A systematic analysis of the transcriptional landscapes associated to PGC1α loss of expression revealed that PGC1α alters the expression of genes encoding for secreted proteins. Cell secretome studies corroborated that PGC1α-dependent ERRα regulation in prostate cancer cells suppresses the growth of tumor cells exposed to their conditioned media. The integration of *in vitro* and *in vivo* secretomics data and genetic perturbation assays revealed spermidine synthase as a transcriptional target of PGC1α and mediator of a paracrine metabolic growth suppressive effect. Moreover, the activity of the regulatory axis PGC1α-ERRα-SRM was reflected in patients and had prognostic value. Altogether, this work provides unprecedented evidence of the non-cell autonomous tumor suppression role of PGC1α, which broadens the view of this co-regulator as a multifactorial tumor suppressor in prostate cancer.

## INTRODUCTION

Prostate cancer (PCa) exhibits the highest incidence among cancer types in men in the European Union (EU) and represents the third cause of death by cancer in the gender (data retrieved from the World Health Organization 2024). Although there are therapies against PCa with a favourable clinical response, 10-15% of patients relapse and are at risk of developing metastatic cancer. The identification of molecular processes relevant in PCa represents a unique opportunity for both the discovery of prognostic biomarkers as well as the design of innovative combinatorial anticancer strategies.

The past two decades of research have provided cancer researchers with extensive molecular data emanating from high throughput studies in large cancer cohorts that can be exploited to discover unprecedented tumor-regulatory processes [1–8]. Inspired by this concept, we identified the association of reduced Peroxisome proliferator-activated receptor gamma coactivator 1-alpha (PGC1α) expression with PCa progression and metastasis, whereby the PGC1α anti-oncogenic activity was selectively mediated by the nuclear receptor ERRα [1, 9, 10]. The PGC1α/ERRα axis suppresses PCa cell proliferation, migration, invasion and metastatic outgrowth, through the regulation of cytoskeleton organisation [11], the elevation of nutrient catabolism [1] and the suppression of polyamine synthesis [12]. Polyamines are polycationic metabolites that are produced from methionine and arginine, and that sustain fundamental cellular processes, such as cell growth and proliferation [13]. Bioactive polyamines predominantly comprise spermidine and spermine that promote key biological activities related to cell growth and proliferation [14, 15]. In addition, polyamines are secreted and can exert paracrine functions [16, 17].

There is increasing evidence supporting a paracrine regulation of cancer cell aggressiveness [18, 19]. Cancer secretomes reprogram the local tumor environment, leading to remodelling of the matrix and the stimulation of pro or anti-tumorigenic phenotypes [3, 20–22]. Due to its inherent potential for diagnosis and prognosis, the deregulation of the secretome composition, at both transcriptomic and proteomic levels, has been a valuable source for the identification of tumor aggressiveness biomarkers in different cancer types, although very little attention has been paid to PCa.

Here we show that the prostate tumor suppressor PGC1α exerts a paracrine growth-inhibitory effect on cancer cells, through the regulation of the secretome composition, and this phenotype is dependent on its transcriptional partner ERRα and restricted to the protein soluble fraction of the secretome. Integrative *in vitro* and *in vivo* secretomics analysis revealed spermidine synthase (SRM) as the common secreted protein whose expression is repressed by the PGC1α-ERRα axis. Mechanistically, we demonstrate that spermidine synthase (SRM) repression is a major contributor to the paracrine PCa suppressive phenotype driven by PGC1α. Moreover, the inverse correlation between SRM and PGC1α expression is reflected in PCa patients. Importantly, monitoring the expression of both genes improve the identification of individuals that will develop aggressive and lethal PCa, opening the window for new therapeutic opportunities based on precision medicine.

## RESULTS

### PGC1α exerts a non-cell autonomous anti-proliferative effect in prostate cancer cells

We previously described the tumor and metastasis suppressive activity of PGC1α-ERRα transcriptional axis in PCa [1, 9]. This complex controls a transcriptional program that goes beyond the regulation of oxidative cell metabolism [1, 10, 11, 23].

In depth gene ontology analysis of the transcriptional landscape associated with PGC1α re-expression in PC3 cells [1] confirmed the increase in the expression of mitochondria-related genes (38.5% of the differential express genes (DEG)), in line with the role of this factor promoting mitochondrial biogenesis [23–25]. Unexpectedly, we observed an enrichment in genes encoding for proteins functionally linked to the extracellular space (Supplementary Figure 1 A), representing more than 26% of the genes transcriptionally deregulated upon PGC1α re-expression. Alterations in the abundance of extracellular or secreted factors are common in different cancer types and can influence disease aggressiveness through paracrine actions [20, 22], although their impact on PCa aggressiveness remains obscure. Therefore, we sought to investigate the possible paracrine effects of PGC1α expression in PCa. As a first approach, we isolated conditioned media (CM) produced by PGC1α non-expressing (PC3 TRIPZ-Pgc1a -Dox) and PGC1α-expressing (PC3 TRIPZ-Pgc1a +Dox) cells, a cellular model in which the transcriptional co-activator is growth-suppressive ([1] and Supplementary Figure 1 B-C). We supplemented different prostate cancer cell lines with these CMs and evaluated their cell number after 7 days of continuous exposure (Figure 1 A-B). Crystal violet staining assays of recipient cells showed that CM produced by PGC1α-positive cells inhibited the growth of PC3, DU145 and 22rv1 cells (Figure 1 B, Supplementary Figure 1 D) with no impact on migration (Supplementary Figure 1 E-F). This anti-proliferative effect of PGC1α-associated CM was dose dependent as increased amounts of CM produced by PGC1α-negative cells abolished the effect (Figure 1 C-D and Supplementary Figure 1 G). Importantly, the observed effects were doxycycline independent, as CM of untreated and doxycycline-treated PC3-TRIPZ cells had no differential impact on the growth of recipient PC3 cells (Supplementary Figure 1 H). Altogether, these data reveal that PGC1α exerts a non-cell autonomous anti-proliferative effect in prostate cancer cells.

**Figure 1.**
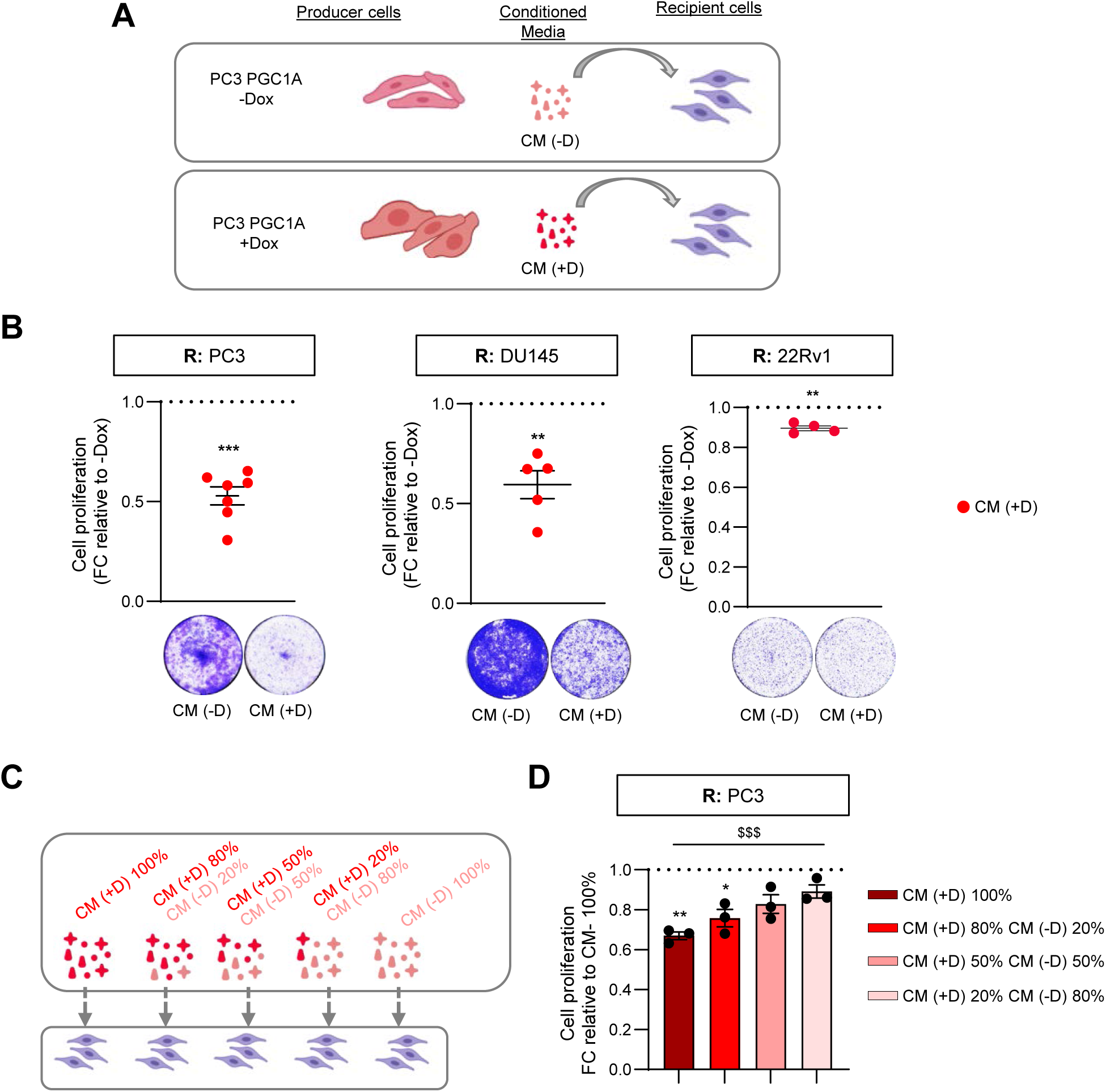
PGC1α-driven conditioned media reduced 2D-proliferation of aggressive prostate cancer cells. **A**. Schematic representation of the experimental approach for condition media production. **B**. Quantification of 2D-cell proliferation in PC3 (n=7; left panel), DU145 (n=5; middle panel) and 22Rv1 (n=4; right panel) cells grown with differential conditioned media produced by PGC1α non-expressing (CM (-D)) and expressing PC3 cells (CM (+D)). A representative image of the crystal violet staining is included below the quantifications. **C**. Schematic representation of the experimental approach for production and combination of conditioned medias. **D**. Dose dependent effect of PGC1α-expressing PC3 cells’ conditioned media (CM (+D)). Different percentages of PGC1α-expressing and non-expressing PC3 conditioned media were used to grow and monitor 2D proliferation of recipient PC3 cells for 7 days (n=3). In B and D, data are normalized to the CM -Dox (non-PGC1α expressing) condition, depicted by a black dotted line. R: recipient cells. CM: conditioned media. D or Dox: doxycyline. FC: fold change. Statistics: one sample t-test with reference value 1 (B and D), ordinary one-way ANOVA (D, depicted with a dollar symbol). * p.value < 0.05; ** p.value < 0.01; ***/$$$ p.value < 0.001. Error bars indicate s.e.m.

### The paracrine growth-suppressive activity of PGC1α is dependent on ERRα and restricted to the non-vesiculated fraction of the conditioned media

The prostate cancer cell-intrinsic tumor suppressive activity of PGC1α largely relies on ERRα [1, 11]. Therefore, we asked whether the growth-inhibitory paracrine activity of the coactivator required the presence of this nuclear receptor. Conditioned media experiments using prostate cancer cells with inducible expression of PGC1α and CRISPR-CAS9-induced deletion of ERRα ([11]; Supplementary Figure 2 A-B) showed that loss of ERRα in producer cells prevented the paracrine action of PGC1α in recipient cells (Figure 2 A).

**Figure 2.**
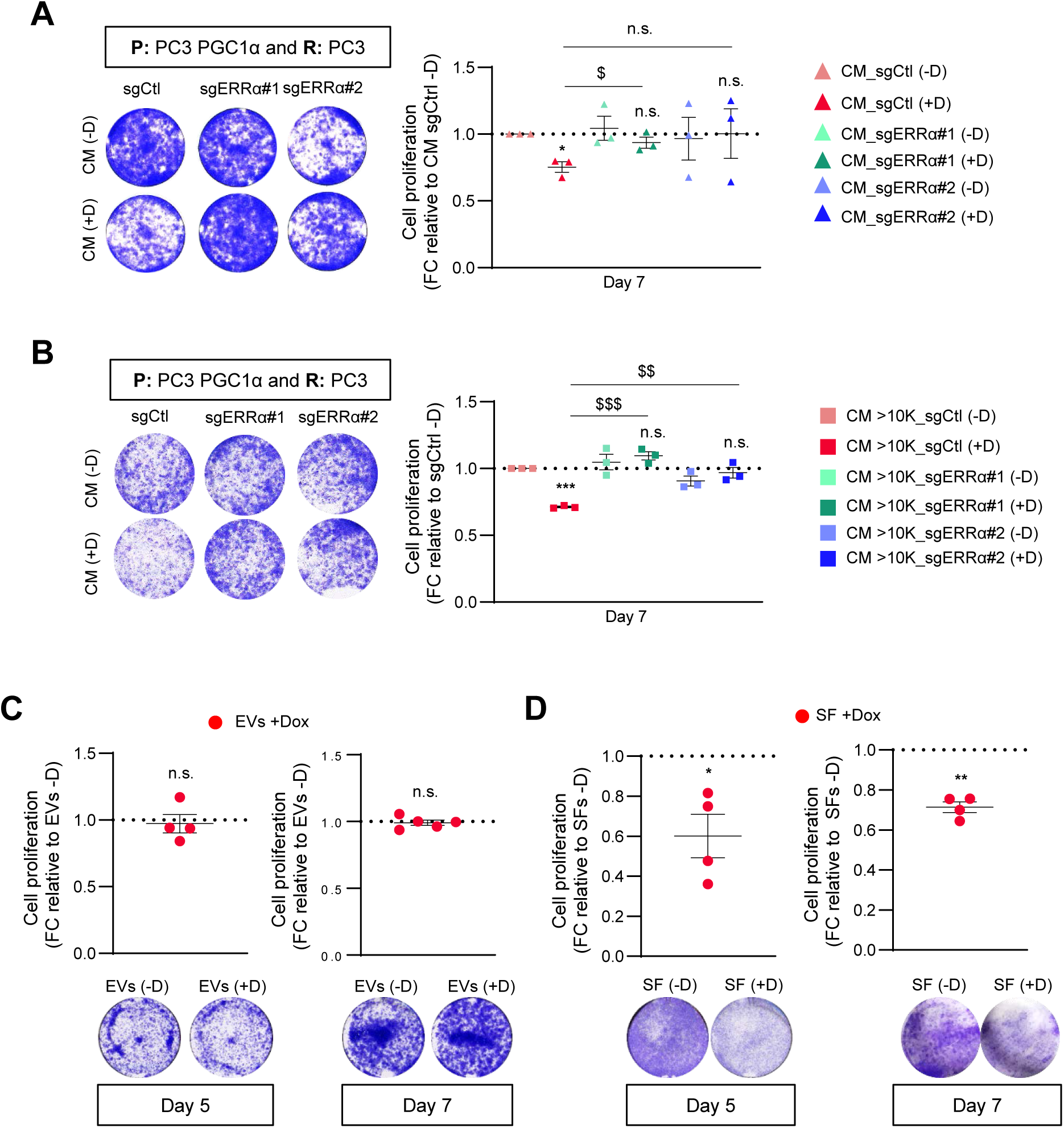
PGC1α- non-cell autonomous anti-proliferative effect is dependent on ERRα and restricted to the soluble fraction of the conditioned media. **A**. Quantification of 2D- cell proliferation (crystal violet) of PC3 (n=3) grown with differential conditioned media produced by PGC1α non-expressing and expressing PC3 cells with or without deletion of ERRα. A representative image of the crystal violet staining is included the quantifications. **B**. Quantification of 2D- cell proliferation (crystal violet) of PC3 (n=3) grown with the heavy fraction of conditioned media (>10 kDa) produced by PGC1α non-expressing and expressing PC3 cells with or without deletion of ERRα. **C**. Effect of extracellular vesicles produced by PGC1α non-expressing and expressing PC3 cells on the 2D- cell proliferation (crystal violet) of PC3 cells during 5 (left panel, n=4) and 7 days (right panel, n=5). **D**. Effect of EVs-depleted fraction produced by PGC1α non-expressing and expressing PC3 cells on the 2D-cell proliferation (crystal violet) of PC3 cells during 5 (left panel, n=4) and 7 days (right panel, n=4). All data are normalized to the CM - Dox (non-PGC1α expressing) condition, depicted by a black dotted line. R: recipient cells. CM: conditioned media. SFs: soluble factors. EVs: extracellular vesicles. D or Dox: doxycyline. Statistics: one sample t-test with reference value 1 (A, B, C and D); paired-t-test (A and B). */$ p.value < 0.05; **/$$ p.value < 0.01; ***/$$$ p.value < 0.001. Asterisks indicate statistical difference between No Dox and Dox conditions and dollar symbols indicate statistical difference between Control Dox and sgERRα#1/sgERRα#2 Dox. Error bars indicate s.e.m.

The factors secreted by cells can be present free in the extracellular media or contained in vesicles [26]. To study which components of the CM were responsible for the paracrine activity of PGC1α in prostate cancer cells, we initially separated the CM into two fractions based on molecular weight: < 10 kDa (light fraction) and > 10 kDa (heavy fraction) and studied their effects on recipient tumor cells. Interestingly, only the heavy fraction preserved the ERRα-dependent growth suppressive activity of PGC1α ectopic expression (Figure 2 B and Supplementary Figure 2 C), thus ruling out a contribution of metabolites and proteins or peptides smaller than 10 kDa.

The heavy fraction of the conditioned media contains proteins heavier than 10kDa as well as extracellular vesicles (EVs). Importantly, EVs play both active and bystander roles in cancer, including PCa [27]. To discriminate between the contribution of EVs and free proteins to the observed phenotype of PGC1α-expressing cells, we separated EVs and the EV-depleted supernatant (Supplementary Figure 2 D and E) and monitored which fraction retained growth-suppressive activity. The uptake of EVs from each producer cell by the recipient PC3 cells was undistinguishable (Supplementary Figure 2 F), and supplementation of culture media with EVs purified from the CM of PC3 PGC1α expressing cells did not suppress prostate cancer cell growth compared to control EVs (Figure 2 C). Importantly, EV-depleted conditioned media from PGC1α-expressing cells exhibited significant growth-suppressive activity (Figure 2 D), suggesting that this effect could be driven by secreted proteins.

### PGC1α regulates the expression and secretion of spermidine synthase in prostate cancer cells

To identify the growth-suppressing secreted factors, we next aimed to characterize the proteomic composition of the PGC1α-associated CMs. Label-free liquid chromatography and mass spectrometry (LC/MS) proteomics analysis revealed 169 differentially abundant secreted proteins in PGC1α-expressing PC3 cells, of them 82 were upregulated and 87 downregulated (Figure 3 A). Functional enrichment analysis of the genes encoding proteins differentially detected in the PGC1α-CM showed an enrichment of extracellular and metabolic proteins (Supplementary Figure 3 A), consistent with the sample type and the canonical metabolic function of PGC1α.

**Figure 3.**
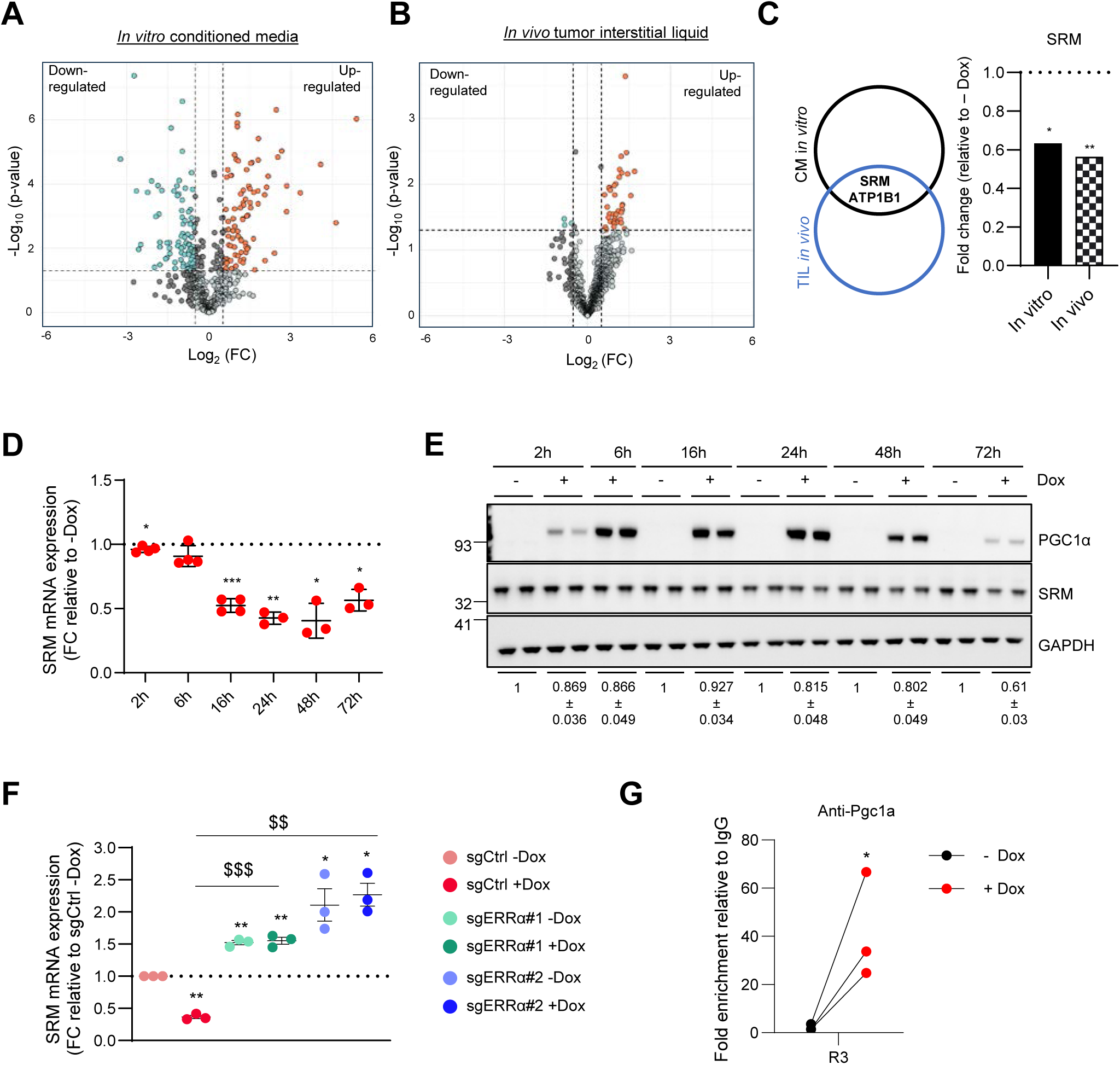
PGC1α regulates spermidine synthase expression and secretion in prostate cancer cells. **A.** Volcano plot representing label-free LC/MS data of proteins differentially secreted by PGC1α expressing and non-expressing PC3 cells. **B**. Volcano plot representing label-free LC/MS data of proteins differentially detected in the tumor interstitial liquid (TIL) isolated from *Pten* and *Pten/Pgc1a* KO prostate tumors. **C**. Venn Diagram (left panel) showing the common secreted proteins differentially detected by LC/MS *in vitro* (CM) and *in vivo* (TIL) and histogram (right panel) showing the degree of change in the detection of SRM. **D-E**. Effect of PGC1α re-expression on SRM in PC3 cells (D, RT-qPCR, n=3; E, one representative Western blot out of 4, quantifications are shown below). **F**. Effect of ERRα deletion on the PGC1α-driven transcriptional deregulation of SRM (RT-qPCR, n=3). **G**. Chromatin immunoprecipitation of exogenous PGC1α on SRM promoter in PC3-PGC1α expressing cells after induction with 0.5 mg/mL doxycycline (n=3). Final data were normalized to IgG (negative immunoprecipitation control). CM: conditioned media. D or Dox: doxycyline. FC: fold change. Statistics: one sample t-test with reference value 1 (D, F, G); paired-t-test (F). */$ p.value < 0.05; **/$$ p.value < 0.01; ***/$$$ p.value < 0.001. Asterisks indicate statistical difference between No Dox and Dox conditions and dollar symbols indicate statistical difference between Control Dox and sgERRα#1/sgERRα#2 Dox. Error bars indicate s.e.m.

To explore the influence of PGC1α on the composition of cell secretomes in a complex biological scenario, we took advantage of our PCa mouse model based on the loss of both *Pten* and *Pgc1α* in prostate epithelia, which leads to invasive carcinoma and metastasis to lymph nodes [1]. We isolated the interstitial liquid of the *Pten* and *Pten/Pgc1α* prostate-conditional knock out tumors and analysed the secretome composition using label free-LC/MS. We detected 41 proteins whose presence in the tumor interstitial liquid (TIL) was altered in double mutant tumors (Figure 3 B), 38 upregulated and 3 downregulated. We then integrated the results obtained *in vitro* and with the murine prostate cancer model and identified two proteins consistently altered upon PGC1α perturbation in PCa, ATPase Na+/K+ Transporting Subunit Beta 1 (ATP1B1) and spermidine synthase (SRM) (Figure 3 C and Supplementary Figure 3 B). ATP1B1 abundance was elevated in PGC1α-expressing cells and *Pten* KO *Pgc1a* WT tumors (Supplementary Figure 3 B), whereas SRM levels were reduced (Figure 3 C) compared to PGC1α non-expressing cells and *Pten/Pgc1a* double KO tumors .

ATP1B1 is a canonical PGC1α-transcriptional target, which we previously reported to be regulated by the coactivator in PCa [1]. However, the lack of scientific evidence on the paracrine role of SRM and its control by PGC1α prompted us to study this candidate.

The tumor suppressive function of PGC1α in PCa is coordinated by transcriptional programs that are driven by ERRα and MYC [1, 11]. Consistently, transcription factor enrichment analysis [28] of the genes encoding for proteins differentially secreted by PGC1α-expressing PCa cells revealed an over-representation of genes canonically regulated by ERRα and MYC (Supplementary Figure 3 C). These data suggested that the differential proteome composition mirrors the cell intrinsic transcriptional reprogramming driven by PGC1α in the producer cells. In line with this notion, time course experiments revealed that SRM mRNA and protein abundance were reduced shortly after PGC1α re-expression (Figure 3 D-E and Supplementary Figure 3 E for protein quantification) independently of doxycycline treatment (Supplementary Figure 3 D). Moreover, this transcriptional regulation was strictly dependent on the presence of ERRα (Figure 3 F).

We sought to deepen in the study of transcriptional regulation of SRM by monitoring the binding of PGC1α to its promoter. We first designed primers that cover the entire SRM promoter based on H3K27Ac open chromatin marks (ENCODE source; Supplementary Figure 3 F) and performed chromatin immunoprecipitation (ChIP) assays. We observed that exogenous PGC1α was bound to the promoter of SRM in a region (R3) that is close to the transcription initiation (Figure 3 G; Supplementary Figure 3 G). Altogether, these data indicate that, in PCa cells, PGC1α negatively and directly regulates the expression of SRM, which is in line with its reduced secretion upon re-expression of the coactivator.

Since polyamine metabolism fuels PCa aggressiveness [29], and some of its metabolic products exhibit paracrine signalling properties [17], we decided to explore the contribution of differential SRM secretion to PCa biology.

#### Reduction in secreted SRM contributes to the paracrine growth-inhibitory action of PGC1α

The alteration in secreted SRM protein levels upon PGC1α expression in PCa cells was suggestive of a contributing function of this enzyme in the control of recipient cell growth. Since SRM produces spermidine and this metabolite is then converted to spermine, we set up ^13^C-labelling metabolic analysis to monitor polyamine biosynthesis in PCa cells supplemented with the CM produced by PGC1α expressing and non-expressing cells. The metabolomics data showed that cells grown in CM derived from PGC1α-positive cells presented reduced levels of spermidine and spermine (Figure 4 A and Supplementary Figure 4 A), consistent with the reported reduction of SRM in the media. We next studied the contribution of SRM to the paracrine suppression of cell growth in PCa cells. On the one hand, we overexpressed SRM in PGC1α-expressing cells to counteract the reduced expression and secretion of this enzyme elicited by the coactivator in PC3 prostate cancer cells (Figure 4 B and Supplementary Figure 4 B). SRM overexpression did not alter cell growth in PGC1α-expressing cells (Supplementary Figure 4 C). However, ectopic expression of this enzyme counteracted the paracrine growth-suppressive effect of the coactivator (Figure 4 C and Supplementary Figure 4 D). On the other hand, we silenced SRM in PC3 cells using two independent constitutive short hairpin RNAs (Figure 4 D and Supplementary Figure 4 E-F) and evaluated the growth-modulatory activity of CM from control and SRM-silenced cells on PC3 recipient cells (Supplementary Figure 4 G). In agreement with our hypothesis, the control CM elicited a greater proliferative response in recipient cells than the one produced by the SRM-silenced counterparts (Figure 4 E).

**Figure 4.**
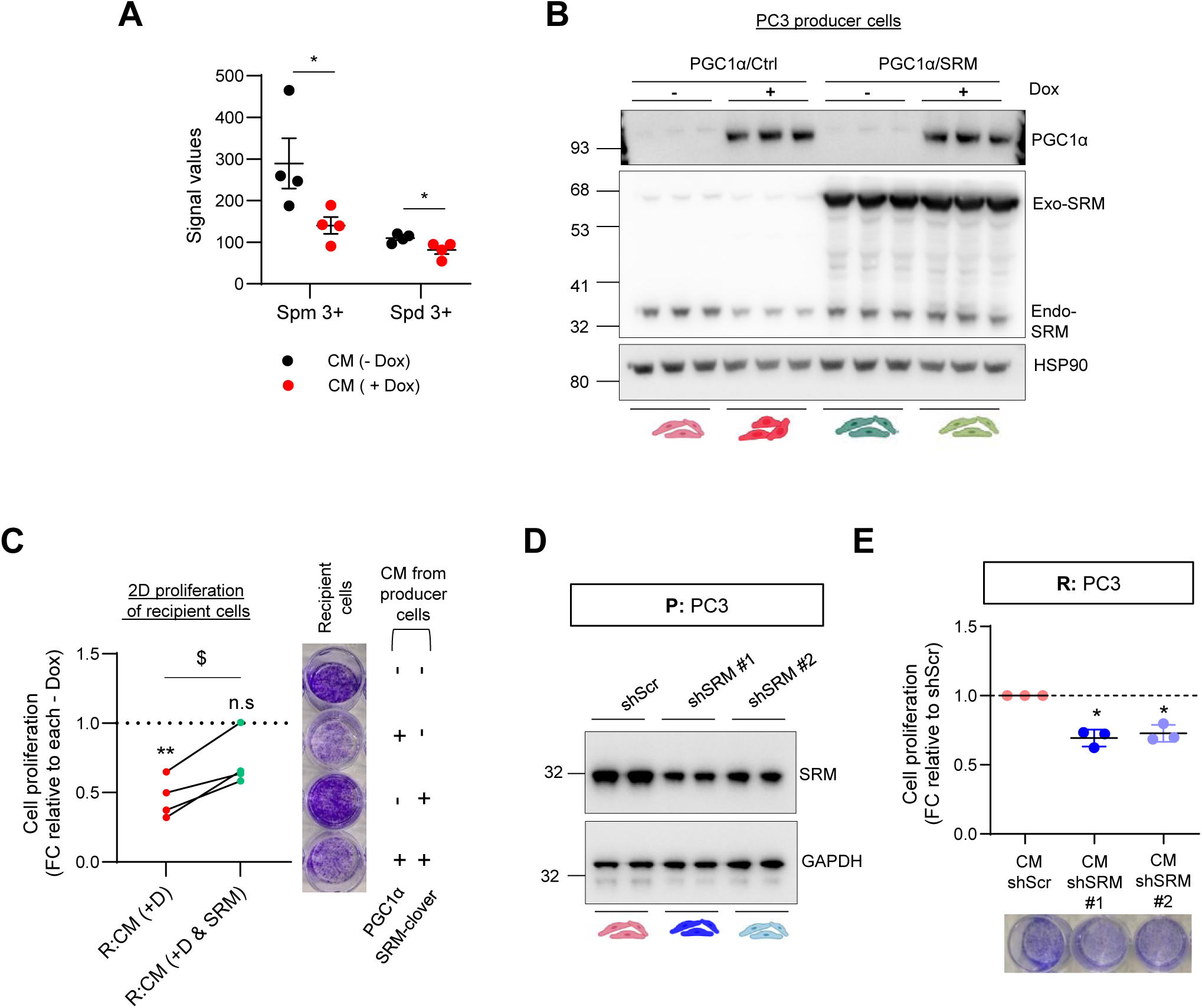
Differential secretion of spermidine synthase contributes to the non-cell autonomous anti-proliferative effect of PGC1α in prostate cancer. **A**. Incorporation of ^13^C from U-13C5-L-Methionine (2h pulse) into spermidine and spermine metabolites after 3 day-treatment of PC3 cells grown with the indicated CM. **B**. Validation of SRM overexpression in PC3 cells with inducible expression of PGC1α (Western blot, one representative image out of 4). **C**. Quantification of 2D-cell proliferation (crystal violet) of PC3 grown with differential CM produced by: PGC1α non-expressing and expressing PC3 cells with or without overexpression of SRM (n=4). **D**. Confirmation of SRM silencing in PC3 cells using two independent short hairpin RNAs (Western blot, one representative image out of 3). **E**. Quantification of 2D-cell proliferation (crystal violet) of PC3 grown with differential CM by PC3 cells in which the expression of SRM was silenced (n=3). In C and E, a representative image of the crystal violet staining is included beside and below the quantifications respectively. In C and E, data are normalized to the CM -Dox (non-PGC1α expressing) condition (C) or to the CM shScr (E), depicted by a black dotted line. P: producer cells. R: recipient cells. CM: conditioned media. D or Dox: doxycyline. FC: fold change. N.s: not significant. Statistics: one sample t-test with reference value 1 (C and E), unpaired-t-test (A, C). Asterisks indicate statistical difference between cells treated with CM produced by PCG1α/Ctrl and PCG1α/SRM cells. */$ p.value < 0.05; ** p.value < 0.01; *** p.value < 0.001. Error bars indicate s.e.m.

Altogether, our data strongly suggest that SRM expression and secretion is under negative control of PGC1α and influences the paracrine communication between cancer cells that further sustains cell growth.

#### The regulation of SRM by PGC1α is conserved in human prostate cancer

We have previously shown that PGC1α expression levels are reduced in PCa and exhibit prognostic potential [1]. Taking advantage of clinically relevant prostate cancer patient cohorts with transcriptomic data ([30–32] and TCGA Firehose Legacy), we monitored the association of SRM mRNA expression with PGC1α transcriptional levels and activity [28]. Consistently with the *in vitro and in vivo* data, prostate tumors from patients with lower expression of PGC1α presented higher levels of SRM mRNA expression (Figure 5 A). In agreement, an inverse correlation between the two genes was also observed when monitoring the continuum of PGC1α expression (Figure 5 B). Moreover, in line with the role of ERRα in the transcriptional effects of PGC1α [1, 11], the analysis of PCa patient transcriptomes revealed a consistent inverse correlation between SRM mRNA levels and the expression of a PCa specific PGC1α-ERRα transcriptional signature [1] (Supplementary Figure 5 A-B).

**Figure 5.**
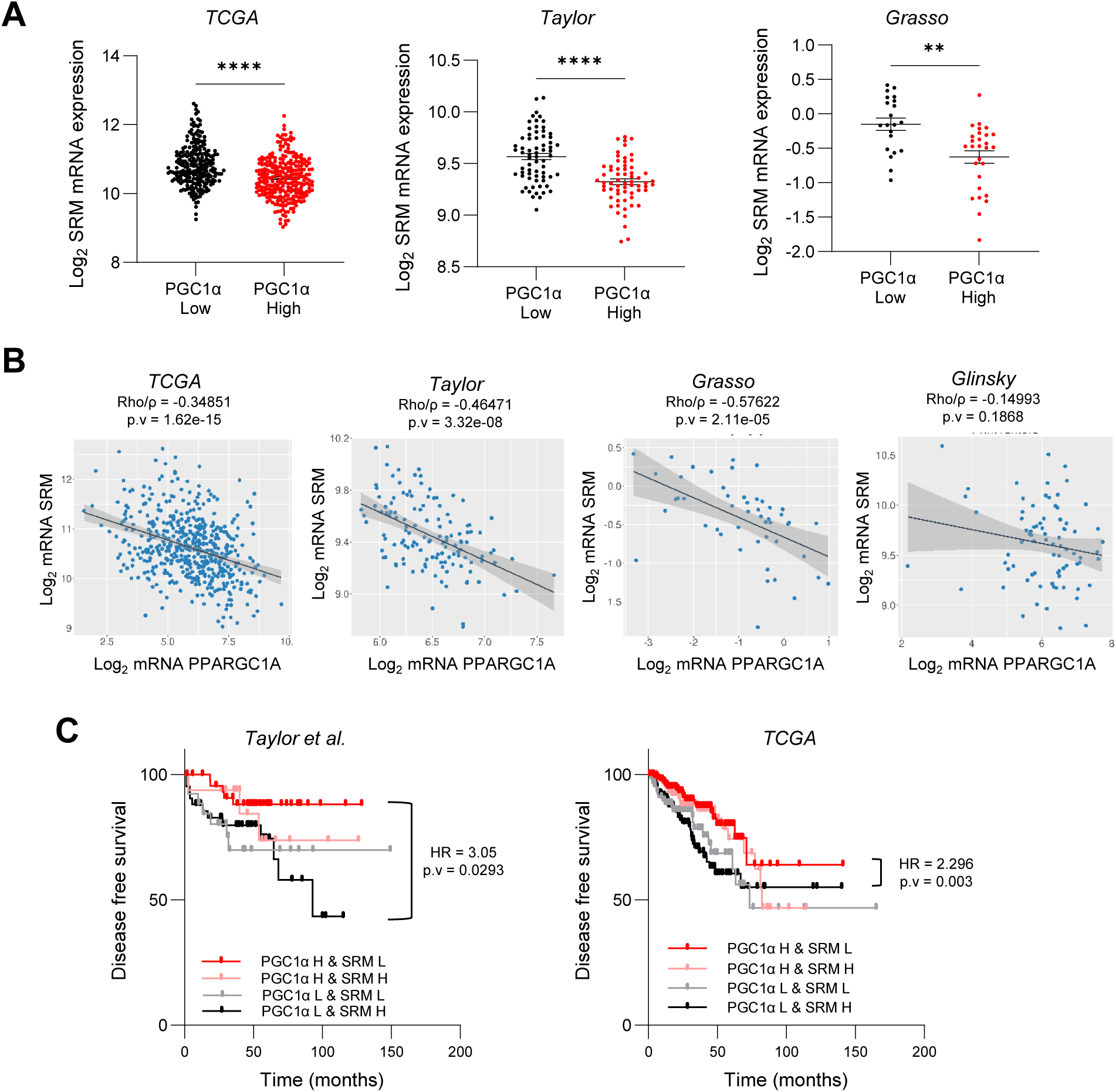
PGC1α expression and activity inversely correlate with SRM in prostate cancer patients and has prognostic value. **A**. Analysis of SRM mRNA in PCa patients stratified according to the mean expression of PGC1α mRNA (PPARGC1A). **B**. Correlation analysis between PPARGC1A and SRM mRNA expression in primary tumor specimens of different prostate cancer datasets. **C**. Association of the combined expression of PPARGC1A and SRM mRNA with disease-free survival (DFS) in human PCa specimens. Patients were grouped according to the average mRNA expression of both genes. H: high, above average. L: low, below average. Four groups were generated: PGC1α H - SRM L, PGC1α L - SRM H, PGC1α H - SRM H and PGC1α L - SRM L. Sample sizes: Grasso, n=45; Taylor, n=131; Glinsky, n=79 and TCGA provisional, n=497. Statistics: Mann Whitney test (A) Spearman correlation Rho/ρ (B), Log-rank (Mantel–Cox) test (C). * p.value < 0.05; ** p.value < 0.01; *** p.value < 0.001. Error bars indicate s.e.m. HR: harzard ratio. p.v: p-value.

We next ascertained whether the newly reported PGC1α-SRM axis could harbour prognostic potential. Using publicly available PCa databases with clinical follow up information matched with transcriptomic data, we classified patients according to the mean expression of SRM and PGC1α mRNAs into SRM or PGC1α Low and High, and generated the different possible combinations. The patient population classified as PGC1A High & SRM Low presented better prognosis than those classified as PGC1A Low & SRM High in two independent cohorts (Figure 5 C).

In conclusion, our study uncovers an unprecedented clinically meaningful paracrine regulation of cell growth governed by the PGC1α-ERRα and elicited, at least in part, by the inhibition of SRM expression and secretion.

## DISCUSSION

Metabolic deregulation is a hallmark of cancer [19] that must be coordinated to contribute to malignant transformation. By proposing transcriptional regulation as a coordination helm driving metabolic rewiring in PCa, in the past we demonstrated the suppressive role of PGC1α [1, 11] although the mechanistic landscape associated to this phenotype is poorly understood. Here we have reported that the transcriptional axis PGC1α-ERRα alters the mRNA expression of genes encoding for secreted proteins, pointing towards a possible non-cell autonomous activity of PGC1α. Secreted factors, through their role as drivers of paracrine cancer cell communication, have previously been described to be active players in therapy resistance and aggressiveness and their transcriptional deregulation contributes to the phenotypic heterogeneity widely observed in cancer patients [20, 22]. In coherence with this suggestion, the integration of transcriptomic data from human (TCGA) and murine models identified dysregulation of secretome genes in PCa [33], although no functional contribution to the disease was assessed.

In our work, we have approached the potential of secreted factors as important contributors of PCa aggressiveness associated to PGC1α dysregulation. The data show that PGC1α exerts a PCa paracrine growth suppressive action that is fully dependent on its transcriptional partner ERRα. Interestingly, this paracrine phenotype is exquisitely led by the protein soluble fraction of PGC1α cell conditioned medias, with no contribution from metabolites or extracellular vesicles (EVs). Beyond its active role in cancer [27, 34], EVs represent a non-invasive tool that may inform about the molecular alterations in PCa [35], therefore we cannot discard the role of PGC1α-associated EVs as surrogate markers of PGC1α activity and therefore PCa status.

Other PCa oncogenic events, such as loss of Pdcd4 or activation of the MNK/eIF4E pathways, have been described to impact cell secretome protein composition affecting and promoting immune evasion and tumor progression [36, 37]. Additionally, PGC1α associated secretomes produced by adipose-derived stem cells have recently been proposed as therapeutic tools against liver fibrosis through the paracrine reduction of human stellate cell proliferation [38]. Therefore, the PCa paracrine suppressive activity that our work has assigned to PGC1α could influence the communication with other cell types and even other acellular components of the tumor microenvironment. Indeed, the functional enrichment analysis of the *in vitro* differential secretomics data showed an enrichment of proteins implicated in extracellular matrix (ECM) production. These data raise new questions on whether PGC1α dysregulation may impact on ECM biology or even on fibroblast function, possibly contributing to PCa aggressiveness.

The paracrine effects of PGC1α described here are specifically retained in the non- vesiculated and proteinaceous fraction of the secretome that is composed, among others, by enzymes. The majority of known secreted enzymes are involved in extracellular matrix degradation and cell migration [34] with very few examples of metabolic enzymes detected in cancer secretomes [39, 40]. Our secretomics data showed an enrichment of metabolic enzymes, some of them known PGC1α targets [1], possibly reflecting the transcriptional status of the CM producer cells. In this work we have shown for the first time, the detection of spermidine synthase (SRM) in the secreted fraction of cancer cells, in both *in vitro* and *in vivo* scenarios. Consistently, we confirmed that changes in SRM secretion were the result of a cell intrinsic transcriptional repression by PGC1α. Highlighting the clinical relevance of this regulatory axis, correlation analysis in PCa patients databases supports this inverse relationship between PGC1α and SRM gene expression.

Although SRM is not a canonical secreted protein (no signal peptide for classical secretion) it is included in the SEPDB database [41–43] therefore it may be secreted by unconventional mechanisms [44]. The data presented in this study not only support this novel localisation of SRM, but also demonstrate its active and novel role in paracrine cell communication. Even though active SRM recombinant protein was not commercially available, we could prove the direct contribution of SRM to the paracrine PCa growth suppression through exogenous genetic rescue of SRM levels and its endogenous silencing. Intriguingly, SRM rescue partially overcomes the paracrine suppressive activity of PGC1α, indicating that additional events may contribute to this novel phenotype assigned to PGC1α in PCa, such as cytokines [45] [46].

The contribution of SRM to the suppressive PGC1α-phenotype was exclusively at the paracrine level (Figure 4C) as the cell intrinsic rescue of SRM in PC3- PGC1α expressing cells does not overcome the growth suppressive phenotype induced by the co-regulator in the CM-producer cells (Supplementary Figure 4 C and [1]). We previously showed that PGC1α re-expression in PCa cells induces a strong cell cycle arrest associated to a profound reduction of MYC expression [11, 12] among several other genes involved in cell proliferation [1]. This strong suppressive phenotype could hardly be rescued by the modulation of a single gene. In line with this idea and in concordance with previous reports [47], we have observed that in *in vitro* full media conditions, the sole SRM perturbation, either overexpression or silencing does not impact on PCa cell proliferation, suggesting that additional intrinsic alterations beyond SRM are required for a full perturbation of cell proliferation in PCa cells. In contrast, under nutrient-poor conditions such as the one induced in CM experiments, cell proliferation is very sensitive to reduced levels of SRM, induced either by PGC1α or by its genetic silencing. Restoring SRM to normal levels provides a clear advantage in these scenarios.

Targeting polyamine metabolism has been proposed as a therapeutic approach in cancer, but these strategies have focused on polyamine depletion through pharmacologically inhibition of enzymes [13, 48]. The data presented herein point towards gene expression inhibition of SRM to reduce paracrine PCa cell growth.

In summary, altogether our data support the notion that cell intrinsic molecular alterations in PCa may play critical roles beyond cell boundaries, expanding our knowledge of the disease and opening windows for new therapeutic opportunities for aggressive PCa.

## METHODS

### Reagents

Doxycycline hyclate (Sigma #D9891) was used to induce gene expression in vectors under tetracycline control. Puromycin (Sigma #P8833) was used for cell selection after lentiviral transfection.

#### Animals

All mouse experiments were carried out following the ethical guidelines established by the Biosafety and Welfare Committee at CIC bioGUNE. The procedures employed were carried out following the recommendations from Association for Assessment and Accreditation of Laboratory Animal Care International. GEMM experiments were generated and carried out as reported in a mixed background [1]. The Pten loxP and Pgc1a loxP conditional knockout alleles have been described elsewhere [59, 60]. Prostate epithelium specific deletion was effected by the Pb-Cre4 [59]. Mice were fasted for 6 h prior to tissue harvest (9 am-3 pm) to prevent metabolic alterations due to immediate food intake.

Tumor interstitial liquid (TIL) was isolated from three-month Pten^pc-/-^ (KO) and Pten^pc-/-^ Ppargc1a^pc-/-^ (DKO) mice. Following the ethical guidelines, mice were sacrificed and the prostate (anterior, ventral and dorso-lateral lobes) was extracted. TIL was obtained through centrifugation for 10 min at 1500 rpm and 4°C. Tissue and TIL were separated and snap-frozen in liquid nitrogen and stored at -80°C for further analysis.

#### Cell culture

Human prostate carcinoma cell lines (PC3 and DU145) were purchased from the Leibniz Institute DSMZ (Deutsche Sammlung von Mikroorganismen und Zellkulturen GmbH) and from the American Type Culture Collection (ATCC), in the case of the 22Rv1 cell line. Both entities provided authentication certificate. PC3 and DU145 cell lines where cultured in Dulbecco’s Modified Eagle Medium without pyruvate (DMEM; Gibco Ref. 41965-039) and 22Rv1 in RPMI (Gibco 61870-010; with GlutaMAX supplement). All of them were cultured with 10% volume for volume (v/v) FBS and 1% (v/v) penicillin– streptomycin and at 37°C in a humidified atmosphere of 5% CO2. All the experiments were performed using this complete media, although, for secretome, soluble factors and EVs isolation experiments, DMEM without pyruvate depleted from bovine derived EVs was prepared. Briefly, 50 ml FBS were diluted in a 1:1 proportion with DMEM without pyruvate. The mixture was ultracentrifuged at 100,000 x g for 16 hours and at 4 °C. Supernatants were poured to the remaining bottle of DMEM without pyruvate and 1% P/S was added. Media (DMEM Exo-free) was filtered through 0.22 µm pores and stored at 4 °C.

Cell lines where periodically subjected to microsatellite-based identity validation. None of the cell lines used in this study were found in the database of commonly misidentified cell lines maintained by the International Cell Line Authentication Committee and NCBI Biosample. 293FT cells were used for lentiviral production. All cell lines were routinely monitored for Mycoplasma contamination. For PGC1α expression, cells were transduced with a modified TRIPZ (Dharmacon) doxycycline-inducible lentiviral construct as previously described [1]. For ESRRA deletion, single-guide RNA (sgRNA) constructs targeting ESRRA (sgERRα#1: 50 CTCCGGCTACCACTATGGTGTGG30; sgERRα#2: 30 AGGAACCCTTTGGACTGTCAGGG50) were designed and cloned as previously described [11]. Two independent lentiviral vectors constitutively expressing validated shRNA against human SRM were obtained from the Mission shRNA Library (TRCN0000290714 and TRCN0000290784). Sequence of human SRM was extracted from pLenti-EFS-FLTID-SRM using EcoRI and NotI restriction sites and subsequently cloned into pCLOVER-RBXN to generate the plasmid pCLOVER-RBXN-SRM. The final construct was verified by DNA sequencing (Eurofins Genomics, Köln, Germany). 293FT cells were transfected with lentiviral vectors and packaging systems following standard procedures, and viral supernatant was used to infect cells. Selection was done using puromycin (2 mg/mL) for 3 days.

#### Conditioned media, extracellular vesicle and soluble factor fraction production and isolation

Due to the previously described anti-proliferative effect of PGC1α in PCa cells [1], the number of PGC1α positive and negative cells was adjusted to have a similar number of producer cells at the day of collection. Therefore, 3x10^6^ and 7x10^6^ cells were plated in PGC1α-negative and positive conditions respectively.

Conditioned media was produced in 150 mm plates and harvested after 48 hours of doxycycline treatment plus additional 24 hours after replacement with fresh media. Briefly, conditioned media was collected and centrifuged at 500 x g, 10 minutes and 10°C to discard cell debris. In parallel, cell number of the producer cells was assessed and a pellet of cells from each condition was taken to ensure the differential protein expression of PGC1α, ERRα and SRM between conditions.

For conditioned media fractionation 10K Amicons (Merck Millipore, Ref. UCF901024) where used to separate and concentrate the secretomes by centrifuging at speeds ranging 1,500-5,000 x g.

EVs and soluble fractions were isolated by ultracentrifugation. Briefly, conditioned media was transferred to a fixed angled 45 Ti or 70 Ti rotor (Beckman Coulter) tubes and centrifuged for 20 minutes at 12,000 x g and 10 °C. The pellet enriched in apoptotic bodies and microvesicles was discarded and the supernatant fraction was poured to a fresh rotor tube and centrifuged 70 minutes at 100,000 x g and 10 °C. Pellets obtained after this step were enriched in EVs, and were resuspended in DPBS 1X into a sole pellet and ultracentrifuged again for 70 minutes, at 10 °C and 100,000 x g. Then, supernatant, corresponding to soluble factor fraction was stored and EVs pellet was resuspended in 100 μl of DPBS 1x for further analysis. For EV staining, the lipid-labelling dye 1,1’- DIOCTADECYL-3,3,3’3’-Tetramethylindocarbocyanine Perchlorate (DilC18(3)) (Thermo Fisher, Ref. D3911) was used. EVs obtained after the first ultracentrifuge step where resuspended in 1 ml of DPBS 1X and 3 µl of the fluorescent dye were added and incubated at room temperature (RT) for 5 minutes. Then, 57 µl of BSA 35% were added and incubated at RT for 1 minute. Next, 18 ml of DPBS 1x were pipetted and samples were ultracentrifuged for 70 minutes at 100,000 x g and 10 °C. Supernatants were removed, pellets resuspended again in 18 ml DPBS 1X and centrifuged for another 70 minutes at 100,000 x g and 10°C. Finally, the supernatants were poured from the tubes and stained EVs pellets were resuspended in 100 μl of DPBS 1X.

#### Electron microscopy

EVs samples were processed at the Spanish National Cancer Research Center (CNIO, Madrid) Electron Microscopy Unit. For negative staining, purified EV fractions were applied onto freshly glow-discharged, carbon-coated, 400-mesh copper EM grids at a concentration of 0.1 mg/ml in a final volume and incubated for 1 minute at RT. The grids were placed consecutively on top of three distinct 50 μl drops of MilliQ water, rinsed gently for 2 seconds, laid on the top of two different 50 μl drops of 1% uranyl acetate (pH = 3), and stained for 1 minute. Finally, the grids were gently side blotted for 5 seconds and air dried. Grid visualization was performed on a Tecnai 12 transmission electron microscope (Thermo Fisher Scientific). Images were recorded at 21,900 nominal magnification with a 4kx4k TemCam-F416 CMOS camera (TVIPS).

#### Cellular assays

Cell number quantification with crystal violet was performed as described in [1]. Recipient cells (PC3, DU145 and 22Rv1) were seeded in 12-well plates (PC3 and DU145: 7,000 cells/well and 22Rv1: 12,000 cells/well). Next day, supernatants were removed and whole conditioned media (1 ml/well), soluble fractions (1 ml/well) or EVs (2-4 µg/well) produced by PGC1a expressing and non-expressing PC3 cells with or without ERRα deletion were pipetted to the wells. This process was repeated every two days, up to day 7. Plates were fixed at different time-points with 10% formalin, washed with 1X PBS and stained with crystal violet [0.1% crystal violet and 20% methanol for 1 hour. Dried crystal violet-stained plates were scanned, and precipitates were dissolved in 10% acetic acid for 30 minutes. Absorbance was measured in 96-well plates in the spectrophotometer (Epoch, Biotek) at a 595 nm wavelength.

For transwell migration assays, PC3 recipient cell lines were treated with conditioned media or EVs obtained from PGC1α-expressing and non-expressing PC3 cell lines. 30,000 PC3 cells were seeded into 6-well plates and underwent conditioned media or EVs treatment during 5 and 6 days respectively. They were then trypsinized, counted and seeded into Boyden chamber transwells (50,000 cells/transwell) resuspended in 500 µl DMEM/well containing 0.5% FBS. Complete culture media (1.4 ml) was pipetted in the bottom well. In parallel, control wells were included as a seeding control of the educated PC3 cells. After 24 hours, migration was stopped: transwells were smoothly cleaned with 1X PBS and, using a cotton bud, the upper side of the transwell membrane was scraped and then rinsed with 1X PBS. Next, transwells were fixed with 10% formalin and stained with crystal violet. Automated inverted Olympus microscope (IX83) (CellSens imaging software) was used to take pictures to further count cell number. Control wells were fixed and stained in parallel to the transwell migration wells. Crystal violet staining was dissolved in 10% acetic acid and absorbance was measured at 595 nm. The values obtained were used to normalize data obtained from the migration assay.

For wound healing assays (WHA), PC3 cells 30.000 cells/ well were seeded into 6-well plates. Twenty-four hours later, media was removed, and cells were treated for five days with conditioned media harvested from PGC1α-expressing and non-expressing PC3 cells. Then, using a 200 µl tip, a longitudinal scratch was performed, supernatants were removed, and fresh differential conditioned media were pipetted. In addition, pictures were taken at the time in which the scratch was performed (time 0 hours). Cells were left to migrate towards the wounded area for 24 hours and pictures were taken at this time point using Olympus Axio Imager A1 CKX3. Data was analysed by means of assessing the area of the initial wound (time 0 hours) minus the area of the wound that remained open after 24 hours of cell migration.

For extracellular vesicle uptake, 200,000 PC3 cells per well were seeded into 6-well plates and left to get attached overnight. Two ml of fresh DMEM Exo-free was added to each well followed by the addition of 2 µg of DilC18(3)-labelled EVs produced by doxycycline-induced and non-induced PC3 TRIPZ PGC1α cells. Three time points (1 hour, 3 hours and 6 hours) were assessed for both conditions, and a negative control of PC3 cells treated with DPBS 1X mixed with DilC18(3) was included. Cells were detached with Cell Dissociation buffer (500 µl/well), centrifuged and pellets were resuspended in 300 µl FACS buffer (PBS 5mM EDTA and 0.1% BSA). Before analysis of EVs uptake using BD Facs Canto devise cell suspensions were passed through CellTrics 50 µm (Sysmex, Ref. 040042-2317).

#### Molecular assays

Western blot was performed as described previously (9). Briefly, cells were seeded on 6-well plates and 4 days after seeding cell lysates were prepared with RIPA buffer (50 mmol/L TrisHCl pH 7.5, 150 mmol/L NaCl, 1 mmol/L EDTA, 0.1% SDS, 1% Nonidet P40, 1% sodium deoxycholate, 1 mmol/L sodium fluoride, 1 mmol/L sodium orthovanadate, 1 mmol/L betaglycerophosphate and protease inhibitor cocktail; Roche).

Protein was quantified using PierceTM BCA Protein Assay Kit (Thermo Fisher Scientific, Ref. 23225). Samples were prepared in Lämmli 5X sample buffer (10% SDS, 50Mm Tris pH 6.8, 10% H2O, 50% Glycerol, 1% β-mercaptoethanol, 0.01M DTT and 0.2 mg/ml of bromophenol blue) and kept at -20 °C for further analysis by western blot. For analysis of EVs by western blot, samples were prepared in non-reducing conditions using Lämmli LDS NuPAGE™ (4X) (Thermo Fisher Scientific, Ref. NP0007).

Protein lysates with Lämmli 1X were boiled at 95 °C for 5 minutes to denaturalize the protein. For EVs samples, boiling was performed at ever increasing temperatures (37 °C, 65 °C and 95 °C), each of them for 5 minutes. Samples were loaded into NuPAGE® Novex® 4-12% Bis-Tris Midi Protein gels (Invitrogen, Ref. NG1403BX10) and run in MOPS SDS buffer (NuPAGE® NP0001-02). For EVs, samples were loaded into Mini-Protean TGX Precast Gels (Biorad, Ref. 456-1085) gels and run in Tris Glycine SDS buffer (National Diagnostics, Ref. EC-870). Both types of gels were resolved at 200 V and transferred to nitrocellulose membranes (Amersham Protran, Ref. 10600001) at 100 V for 1 hour. Membranes were then blocked in 5% non-fat milk prepared in Tris-buffered saline solution containing 0.01% Tween-20 (TBS-T).The following antibodies were used: PGC1α H300 (Santa Cruz Biotechnology #sc-13067), ERRα (Cell Signaling Technology#13826), β-actin (Cell SignalingTechnology #3700S), GAPDH (Cell Signaling Technology #2118), SRM (Proteintech #19858-1-AP), CD9 (R&D Systems #209306), CD63 (Developmental Studies Hybridoma Bank H5C6), GRP78 (BD Biosciences 40/BiP), COX IV (Cell Signaling Technology #11967). All primary antibodies were used at a 1:1,000 dilution, except β-actin (1:2,000). Mouse and rabbit secondary antibodies were purchased from Jackson ImmunoResearch. After standard SDS-PAGE and Western blotting techniques, proteins were visualized using the ECL system in the iBright FL1000 Imaging System and BioRad.

RNA from human prostate cancer cell lines was extracted using NucleoSpin® RNA isolation kit from Macherey-Nagel (Ref: 740955.240C), following the manufacturer’s protocol. RNA concentration was determined using Nanodrop ND-1000 Spectrophotometer. 1 µg of total RNA was used for cDNA synthesis using Maxima H Minus cDNA synthesis with dsDNase. Thermo Scientific, Ref: M1682. Quantitative real-time PCR (qRT-PCR) was performed as described previously [1] and using a QS5 Real-Time PCR System (Applied Biosystems). For detection of SRM gene expression, we used PrimeTimeTM (Integrated DNA Technologies- IDT) TaqMan probe with reference Hs.PT.58.19689793. qRT-PCR data were normalized using GAPDH Hs.PT.39a.22214836 from IDT.

#### Chromatin immunoprecipitation

Chromatin immunoprecipitation (ChIP) was performed using the SimpleChIP Enzymatic Chromatin IP Kit (catalog no. 9003, Cell Signaling Technology, Inc). Four million PC3 TRIPZ-Pgc1a cells per immunoprecipitation were grown in 150-mm dishes either with or without 0.5-μg/mL doxycycline for 16 hours. Cells were cross-linked with 37% formaldehyde for 10 minutes at room temperature. Glycine was added to dishes and cells were incubated for 5 minutes at room temperature. Cells were then washed twice with ice-cold PBS and scraped into PBS þ PIC. Pelleted cells were lysed and nuclei were harvested following the manufacturer’s instructions. Nuclear lysates were digested with micrococcal nuclease for 20 minutes at 37°C and then sonicated in 500-µL aliquots on ice for six pulses of 20 seconds using a Branson sonicator. Cells were held on ice for at least 20 seconds between sonications. Lysates were clarified at 9,400 x g for 10 minutes at 4°C, and chromatin was stored at -80°C. HA-Tag polyclonal antibody (Cell Signaling Technology #3724) and IgG antibody (Cell Signaling Technology #2729) were incubated overnight (4°C) with rotation and protein G magnetic beads were incubated for 2 hours (4°C). Washes and elution of chromatin were performed following manufacturer’s instructions. DNA quantification was carried out using a QS5 Real-Time PCR System (Applied Biosystems) with SYBR Green reagents and primers (shown in Supplementary Table 1) that amplify the regulatory region of SRM promoter based on H3K27Ac marks.

#### Label-Free Proteomic analysis

PC3 TRIPZ PGC1α cells were pre-induced with doxycycline for three days and seeded at high confluences in 100 mm plates (4x10^6^ PGC1α-expressing and non-expressing cells). 24 hours later, supernatants were removed, cells were washed three times with DPBS 1X to remove FBS and serum-free DMEM was added. Three hours later, conditioned media were collected and centrifuged at 500 x g for 10 minutes at 10 °C. Samples were precipitated using the GE Health 2-D Clean-Up Kit (Sigma-Aldrich, Ref. 80-6484-51). Proteins were extracted using 7M urea, 2M thiourea, 4% CHAPS. Samples were incubated for 30 minutes at RT under agitation and digested following the filter-aided sample preparation (FASP) protocol described by Wisniewski and colleagues (Wiśniewski et al. 2009). Trypsin was added to a trypsin:protein ratio of 1:10, and the mixture was incubated overnight at 37°C, dried out in a RVC2 25 speedvac concentrator (Christ), and re-suspended in 0.1% formic acid (FA).

Processed samples were either analysed in an Orbitrap XL ETD mass spectrometer (Thermo-Fisher) or a timsTOF Pro with PASEF (Bruker Daltonics). The Orbitrap XL ETD mass spectrometer was connected to a nanoACQUITY UPLC System (Waters). Sample was loaded onto a Symmetry 300 C18 UPLC Trap column (180 μm x 20 mm, 5 μm (Waters) and resolved in a BEH130 C18 column (75 μm x 200 mm, 1.7 μm (Waters). The mass spectrometer automatically switched between MS and MS/MS acquisition in DDA mode, in an alternating fashion. Full MS survey spectra (m/z 400–2000) were acquired in the Orbitrap with 30000 resolution at m/z 400. The six most intense ions were subjected to CID fragmentation in the linear ion trap. Precursors with charge states of 2 and 3 were specifically selected for fragmentation. Analysed ions were excluded from further analysis for 30 seconds using dynamic exclusion lists. The timsTOF Pro with PASEF was coupled online to a nanoElute liquid chromatograph (Bruker). Sample (200 ng) was directly loaded in a 15 cm Bruker nanoelute FIFTEEN C18 analytical column (Bruker) and resolved at 400 nl/min with a 30-minute gradient. Column was heated to 50°C using an oven.

#### Data analysis and statistics

For proteomics data analysis, Progenesis LC-MS software (Nonlinear Dynamics Ltd., Newcastle upon Tyne, UK) was used for the Orbitrap data. Searches were carried out suing Mascot (Matrix Science). Tolerances of 10ppm and 0.5 Da were used for precursor and fragment searches, respectively. Only peptides passing the FDR < 1% filter were considered for further analysis. Protein quantitation was performed using the information concerning to the three most intense peptides (when available), and only proteins quantified with least two peptides at an FDR<1% were considered for further analysis. On the other hand, data coming from the timsTOF Pro with PASEF was analysed using PEAKS software (Bioinformatics solutions). Searches were carried out against a database consisting of Homo sapiens entries (Uniprot/ Swissprot), with precursor and fragment tolerances of 20 ppm and 0.05 Da. Only proteins identified with at least two peptides at FDR<1% were considered for further analysis. Protein abundances were normalized against the control (no dox) condition per dataset and replicate, loaded onto Perseus platform [61] and further processed (log2 transformation, imputation). A t-test was applied to determine the statistical significance of the differences detected between the corresponding groups.

The differential gene expression analysis driven by PGC1α in PC3 cells [1] can be obtained from GEO with reference GSE75193.

No statistical method was used to predetermine sample size. The experiments were not randomized. The investigators were not blinded to allocation during experiments and outcome assessment. n values represent the number of independent experiments performed, the number of individual mice, or patient specimens. For each independent *in vitro* experiment, normal distribution was assumed, and one-sample t test was applied for one-component comparisons with control and Student t test for two-component comparisons. For *in vivo* experiments, a non-parametric Mann–Whitney exact test was used. Two-tailed statistical analysis was applied for experimental design without predicted result, and one-tailed for validation or hypothesis-driven experiments. The confidence level used for all the statistical analyses was of 95% (alpha value ¼ 0.05). GraphPad Prism 10 software was used for statistical calculations.

Bioinformatic analysis containing gene expression prostate cancer patient data (correlation and gene enrichment analysis) was performed using the web-based interface Cancertool [28]. To determine the correlation between SRM and PGC1α-ERRα gene signature [1], we calculated the value of the signature per individual by comparing the average expression levels of the scaled values of all the genes. For correlation analysis, we applied Spearman correlation (rho) on these values in patient samples using cor.test function in R language.

## Acknowledgments

Apologies to those whose related publications were not cited because of space limitations. We are grateful to the Torrano and Carracedo labs for valuable input. AS, MF and MP were funded by a Basque Government predoctoral grants, AZL is supported by Fundación Científica AECC (INVES223210ZABA), IH was funded by the Juan de la Cierva Incorporación program (IJC2019-040709-I), AP was funded by a predoctoral grant from the UPV/EHU, AM was funded by a FPI predoctoral fellowship from MINECO, LVJ was funded by the Juan de la Cierva Incorporación program (IJC2020-044958-I) and Fundación CRIS Contra el Cáncer PR_TPD_2022-04 in partnership with Fundación Adey, HP was funded by the European Union’s Horizon 2020 research and innovation programme under the Marie Skłodowska-Curie ITN proEVLifeCycle with grant agreement No 860303, EB is supported by the Basque Department of Industry, Tourism and Trade (Elkartek), the MICINN PID2022-141556OB-I00 (FEDER/EU), Fundación Domingo Martínez, and the Severo Ochoa Excellence Accreditation (CEX2021-001136- S funded by MICIU/AEI/10.13039/50110001103), JDS was funded by MCIN/AEI/10.13039/501100011033 projects PID2023-147399NB-I00, PID2020-114178GB-I00, the Severo Ochoa Excellence Program CEX2021-001136-S and CEX2021-001202; FR and JMFP were supported by Project PID2021-125104OB-I00., AC is supported by the Basque Department of Industry, Tourism and Trade (Elkartek), the BBVA foundation (Becas Leonardo), the MICINN (PID2022-141553OB-I0 (FEDER/EU); Fundación Cris Contra el Cáncer (PR_EX_2021-22), Severo Ochoa Excellence Accreditation (CEX2021-001136-S), European Training Networks Project (H2020-MSCA-ITN-308 2016 721532), the AECC (GCTRA18006CARR), Fundación Jesús Serra, iDIFFER network of Excellence (RED2022-134792-T), and the European Research Council (Consolidator Grant 819242). The work of V Torrano was supported by Ramón y Cajal Program RYC-2017-22295, national grants RTI2018-097267-B-I00 and PID2021-123372OB-I00 from MINECO, Fundación Científica AECC LABAE211656TORR, the Basque Department of Industry, Tourism and Trade (ELKARTEK24/10), the Department of Education IKERTALDE IT1720-22, Consolidación Investigadora 2023 CNS2023-143848, Fundación FERO, ĹOreal For Women in Science. CIBERONC was cofunded with FEDER funds and funded by ISCIII.

## Author contributions

Conception and design: Schaub-Clarigué A and Torrano V.

Development of methodology: Schaub-Clarigué A, I Hermanova and Torrano V

Acquisition of data (provided animals, acquired and managed patients, provided facilities, etc.): Schaub-Clarigué A, Hermanova I, Pintor A, Macchia A, Valcarcel-Jimenez L, Fagoaga M, Zabala-Letona A, Pujana-Vaquerizo M, Royo F, Peinado H, Azkargorta M, Elortza F, Falcón-Pérez JM, Carracedo A and Torrano V

Analysis and interpretation of data (e.g., statistical analysis, biostatistics, computational analysis): Schaub-Clarigué A, Hermanova I, Fagoaga M, Garcia-Longarte S, Azkargorta M, Elortza F, Berra E, and Torrano V

Writing, review, and/or revision of the manuscript: Torrano V wrote the manuscript and all the rest of the authors were involved in its revision.

Administrative, technical, or material support (i.e., reporting or organizing data, constructing databases): Lectez B, Sutherland J, Garcia-Longarte S

Study supervision: Torrano V

## Notes

### Competing Interest Statement

The authors have declared no competing interest.

