## Supplementary Figures for "Secreted spermidine synthase reveals a paracrine role for PGC1α-induced growth suppression in prostate cancer"

### Slide 1
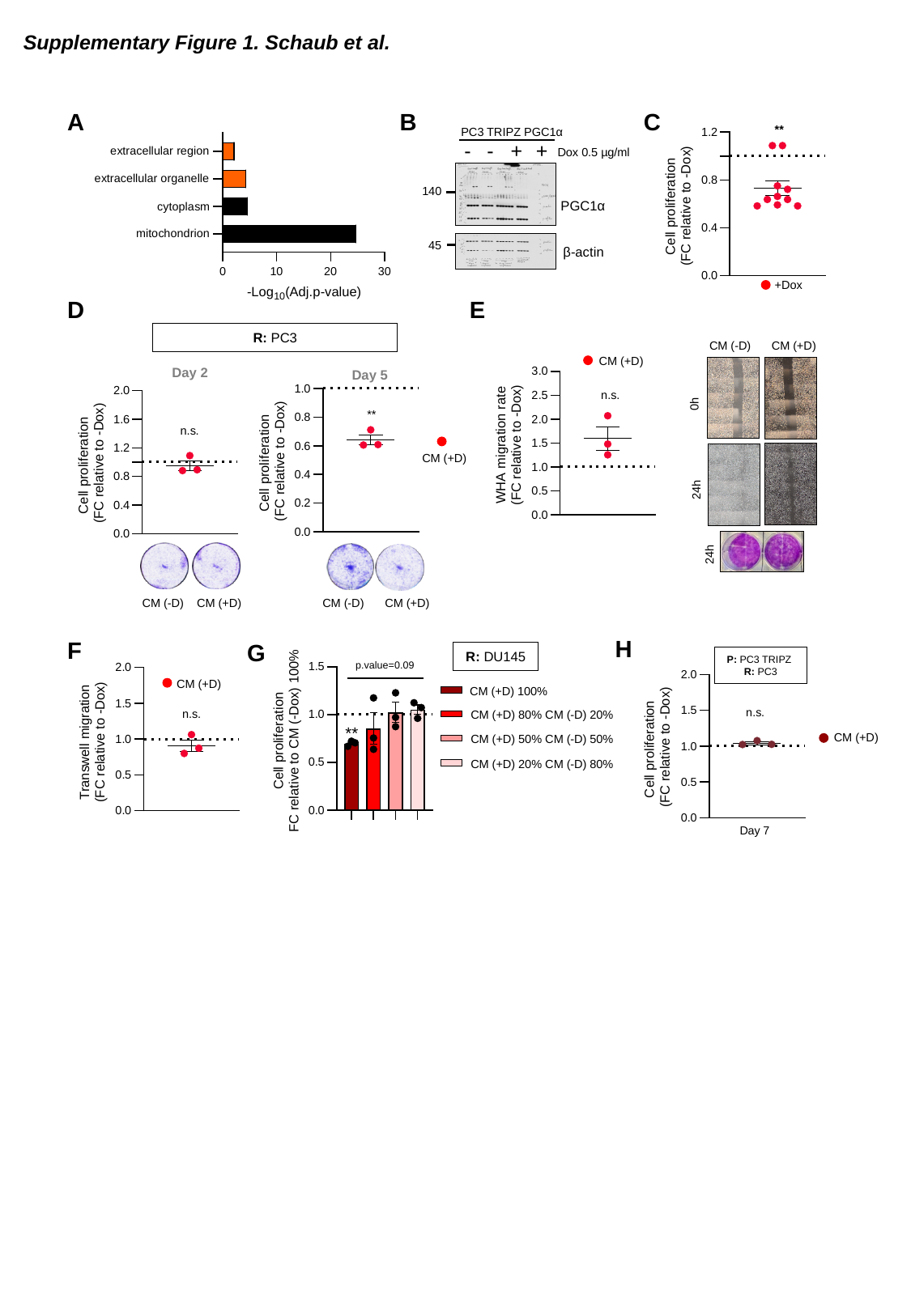

Supplementary Figure 1. Schaub et al.
A
B
C
PC3 TRIPZ PGC1α
+
-
-
+
Dox 0.5 µg/ml
140
PGC1α
45
β-actin
+Dox
E
D
R: PC3
CM (-D)
CM (+D)
CM (+D)
Day 2
Day 5
0h
CM (+D)
24h
24h
CM (-D)
CM (+D)
CM (-D)
CM (+D)
H
F
G
R: DU145
P: PC3 TRIPZ
R: PC3
CM (+D)
CM (+D) 100%
CM (+D) 80% CM (-D) 20%
CM (+D)
CM (+D) 50% CM (-D) 50%
CM (+D) 20% CM (-D) 80%

### Slide 2
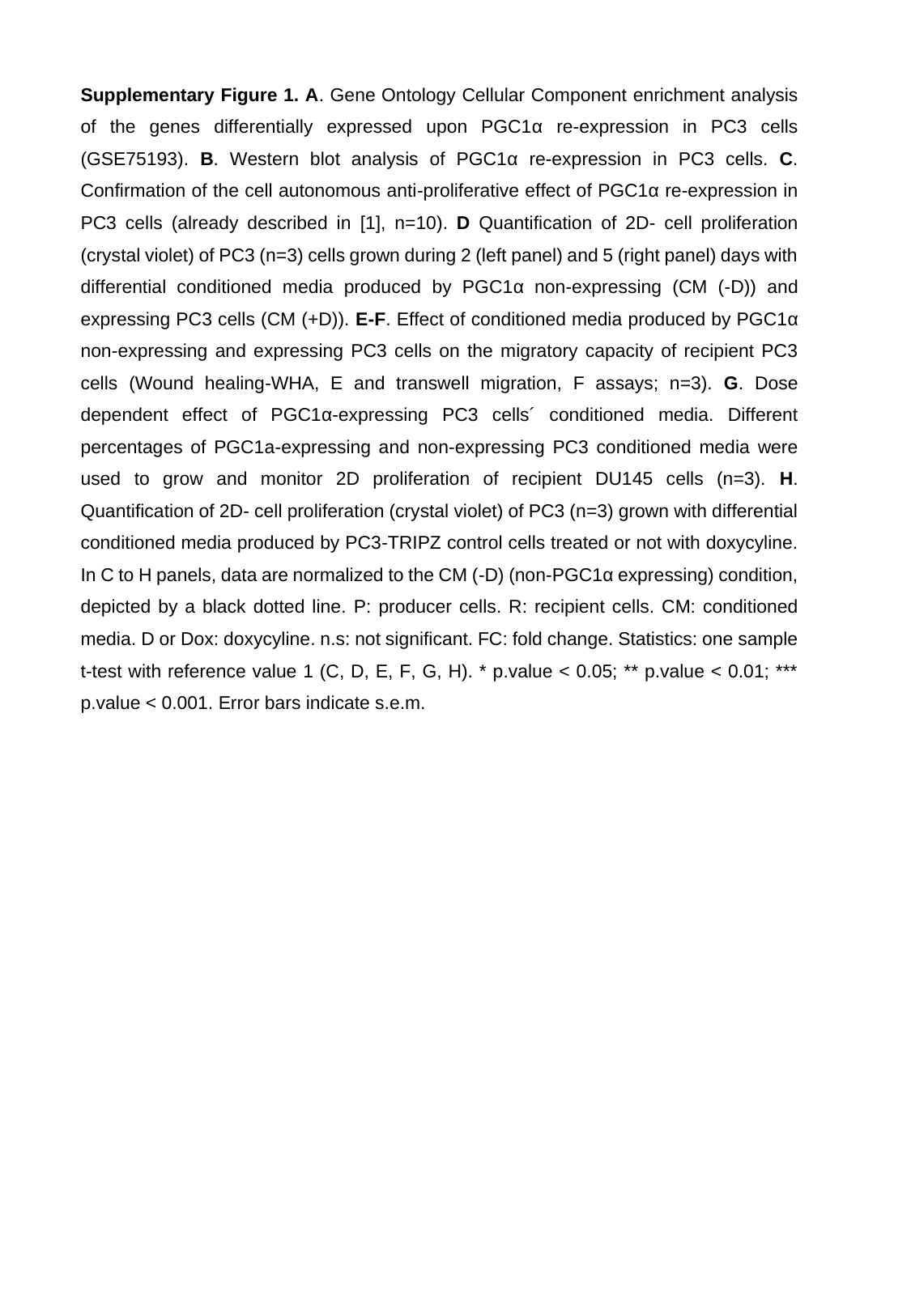

### Slide 3
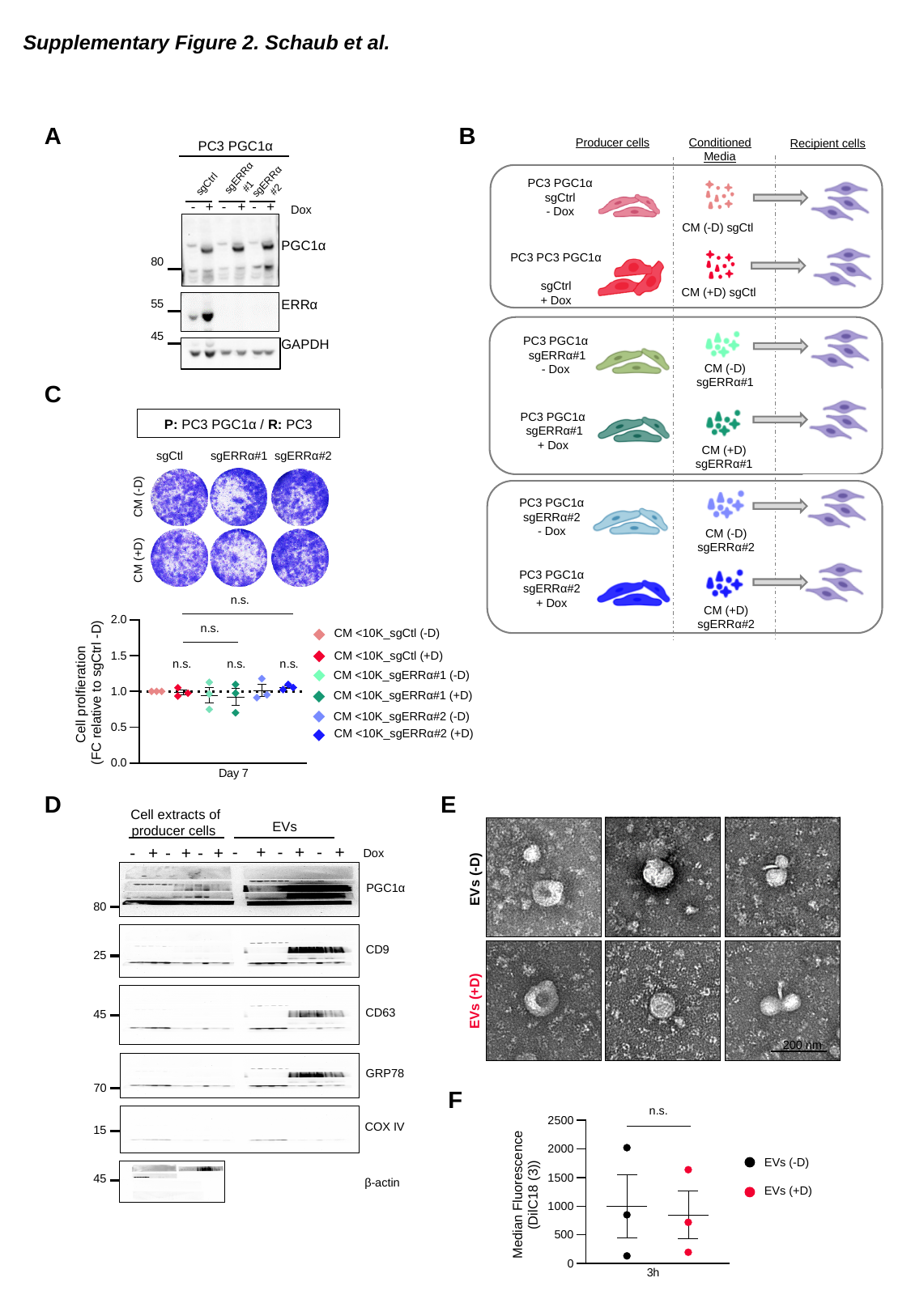

Supplementary Figure 2. Schaub et al.
B
A
Producer cells
Conditioned Media
PC3 PGC1α
Recipient cells
sgERRα
#1
sgERRα
#2
sgCtrl
PC3 PGC1α
sgCtrl
- Dox
-
+
-
+
-
+
Dox
CM (-D) sgCtl
PGC1α
80
PC3 PC3 PGC1α
sgCtrl
+ Dox
CM (+D) sgCtl
ERRα
55
45
GAPDH
PC3 PGC1α
 sgERRα#1
- Dox
CM (-D) sgERRα#1
C
P: PC3 PGC1α / R: PC3
PC3 PGC1α
 sgERRα#1
+ Dox
CM (+D) sgERRα#1
sgCtl
sgERRα#1
sgERRα#2
CM (-D)
PC3 PGC1α
sgERRα#2
- Dox
CM (-D) sgERRα#2
CM (+D)
PC3 PGC1α
sgERRα#2
+ Dox
CM (+D) sgERRα#2
CM <10K_sgCtl (-D)
CM <10K_sgCtl (+D)
CM <10K_sgERRα#1 (-D)
CM <10K_sgERRα#1 (+D)
CM <10K_sgERRα#2 (-D)
CM <10K_sgERRα#2 (+D)
D
E
 Cell extracts of
producer cells
 EVs
-
+
-
+
-
+
-
+
-
+
-
+
Dox
EVs (-D)
PGC1α
80
CD9
25
EVs (+D)
CD63
45
200 nm
GRP78
70
F
COX IV
15
EVs (-D)
45
β-actin
EVs (+D)

### Slide 4
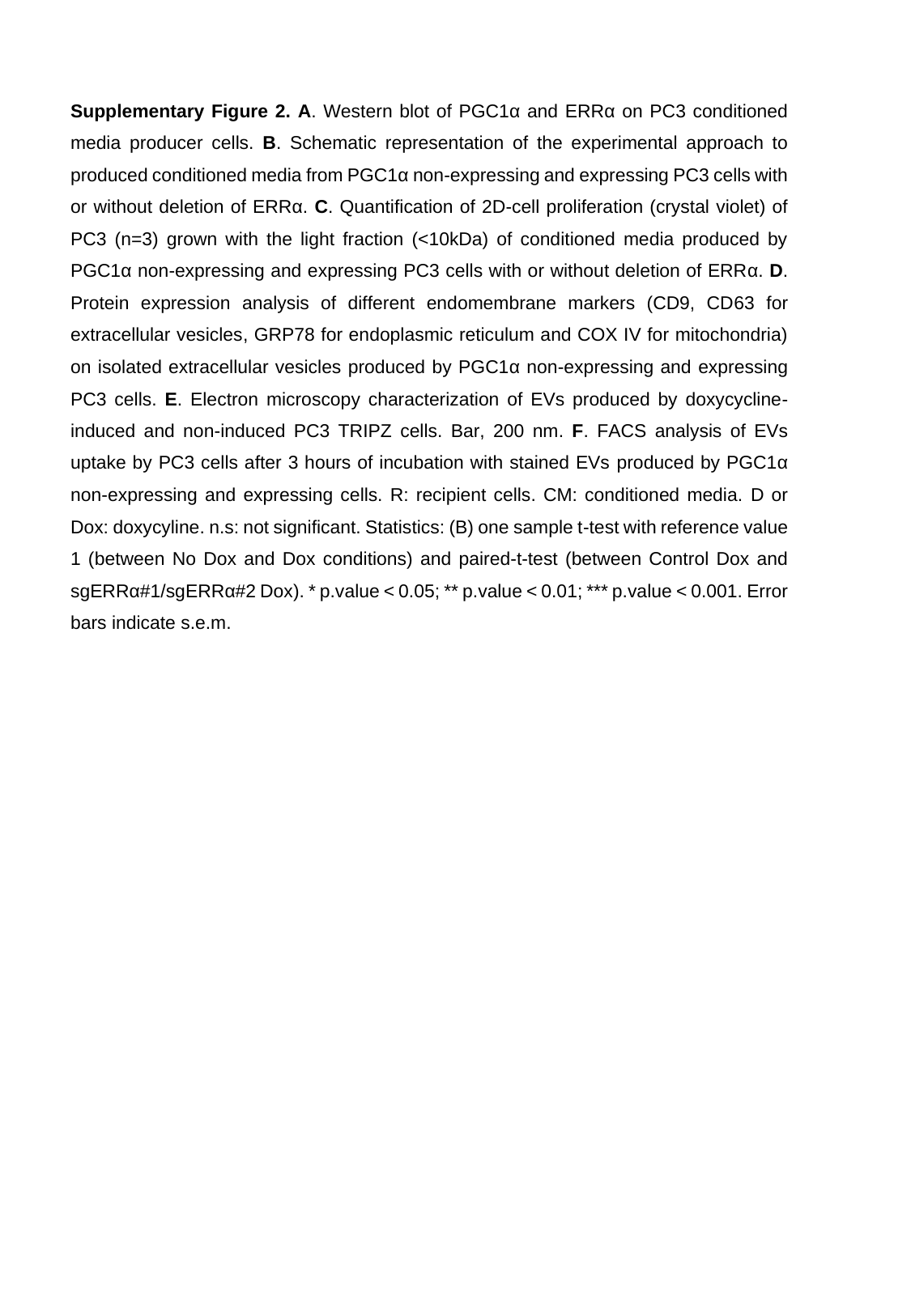

### Slide 5
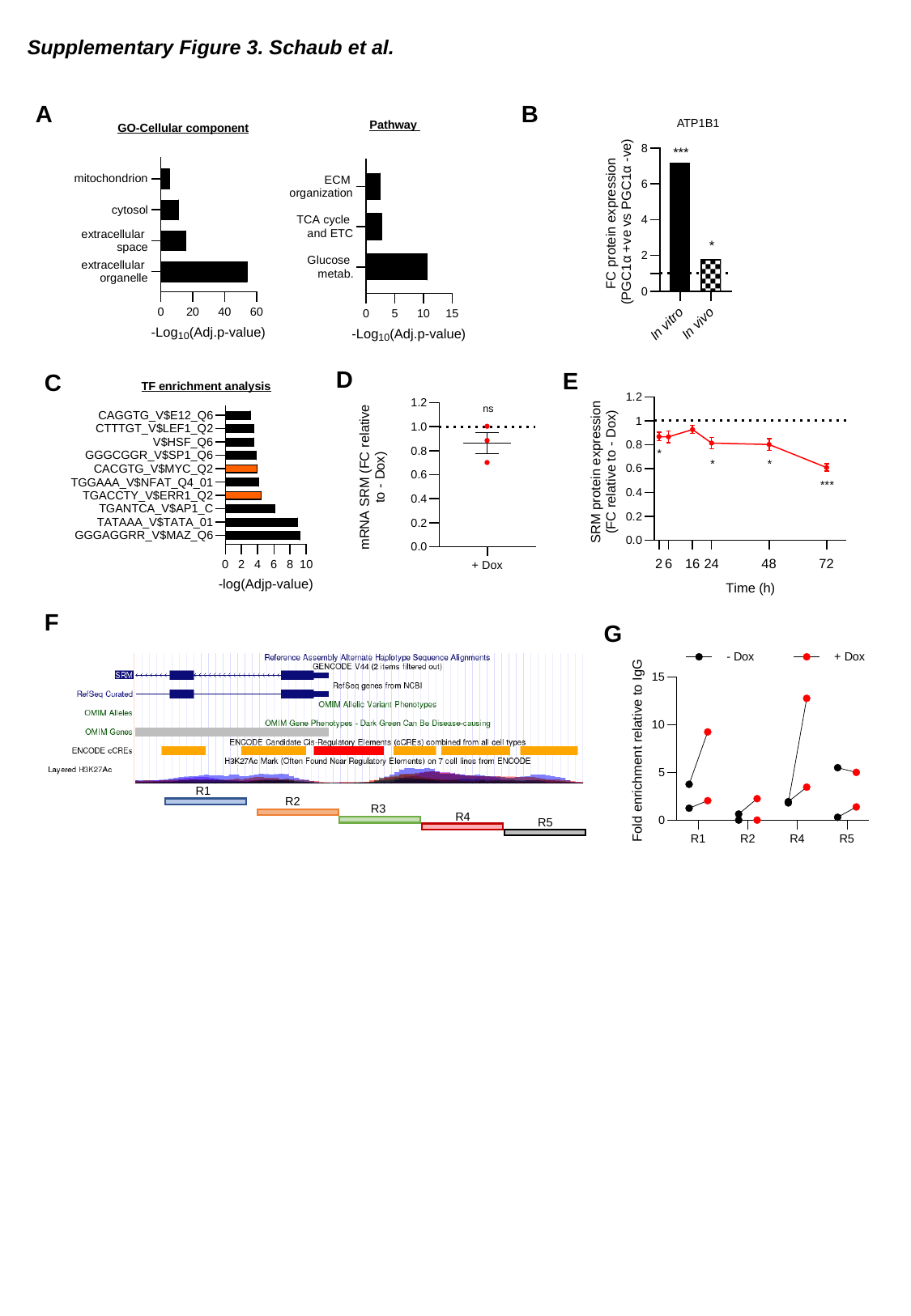

Supplementary Figure 3. Schaub et al.
A
B
ATP1B1
Pathway
GO-Cellular component
***
*
D
E
C
TF enrichment analysis
F
G
R1
R2
R3
R4
R5

### Slide 6
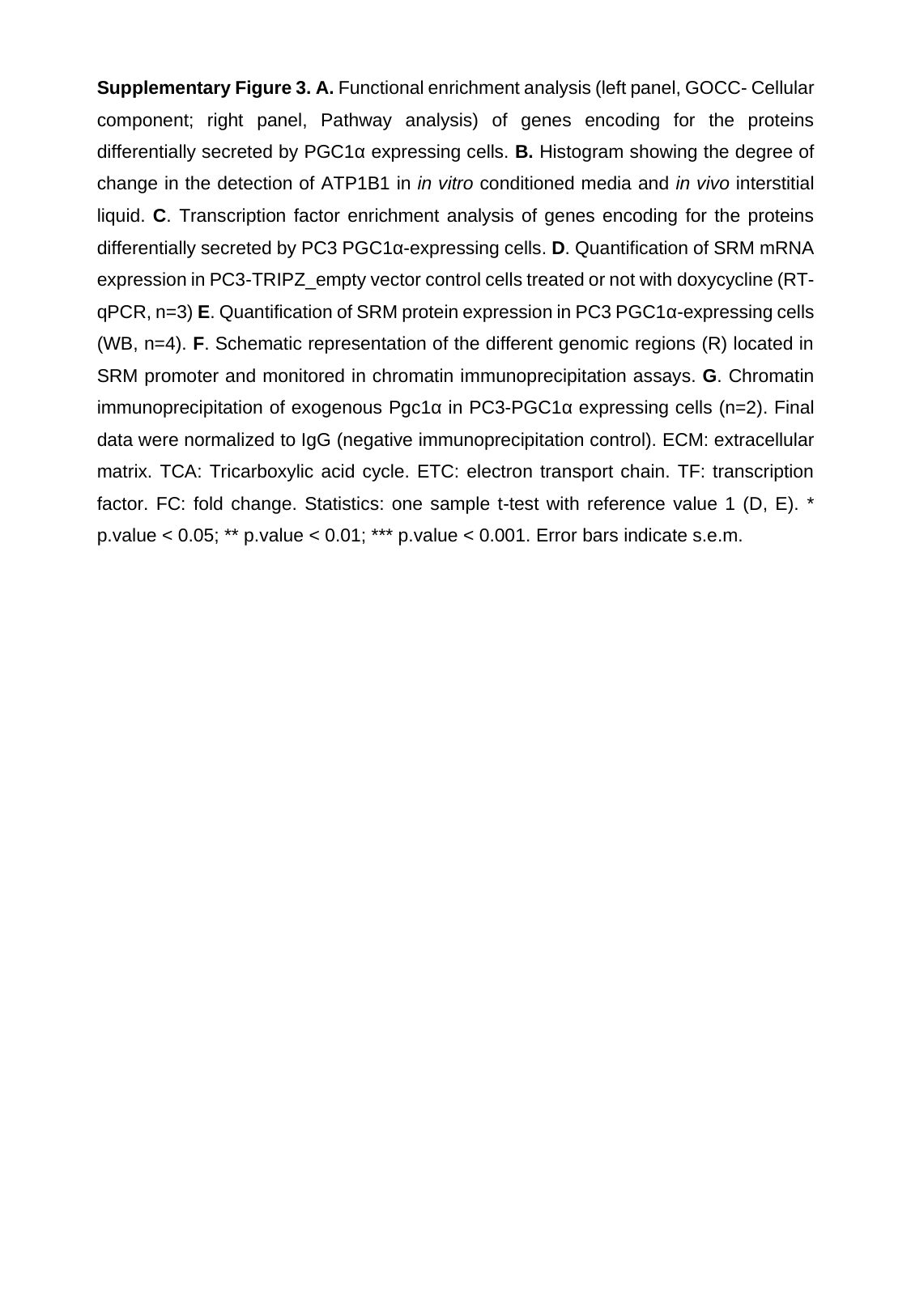

### Slide 7
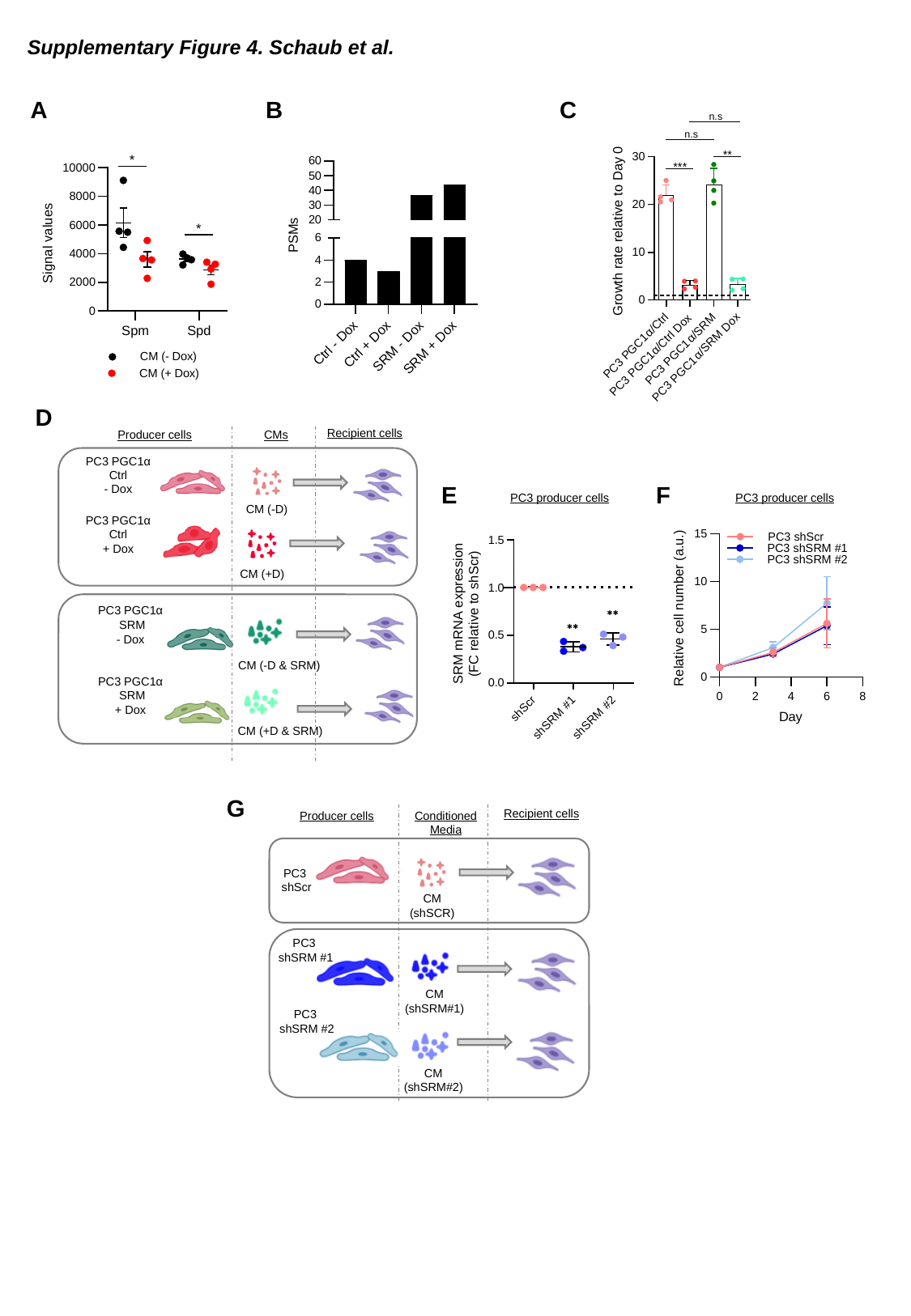

Supplementary Figure 4. Schaub et al.
A
B
C
*
*
D
Recipient cells
Producer cells
CMs
PC3 PGC1α
Ctrl
- Dox
CM (-D)
PC3 PGC1α
Ctrl
+ Dox
CM (+D)
PC3 PGC1α
 SRM
- Dox
CM (-D & SRM)
PC3 PGC1α
 SRM
+ Dox
CM (+D & SRM)
E
F
PC3 producer cells
PC3 producer cells
G
Producer cells
Conditioned Media
Recipient cells
PC3
shScr
CM (shSCR)
PC3
shSRM #1
CM (shSRM#1)
PC3
shSRM #2
CM (shSRM#2)

### Slide 8
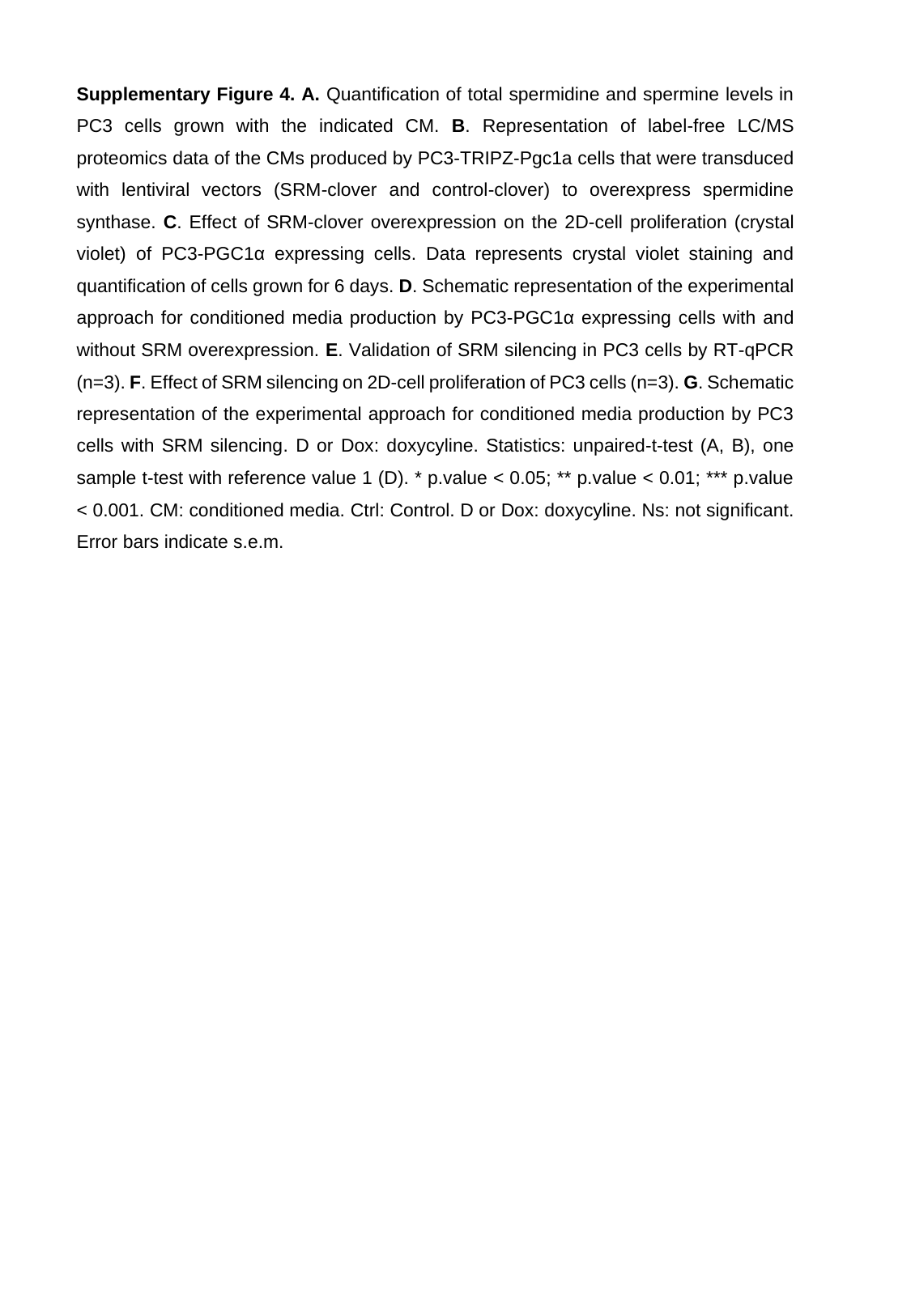

### Slide 9
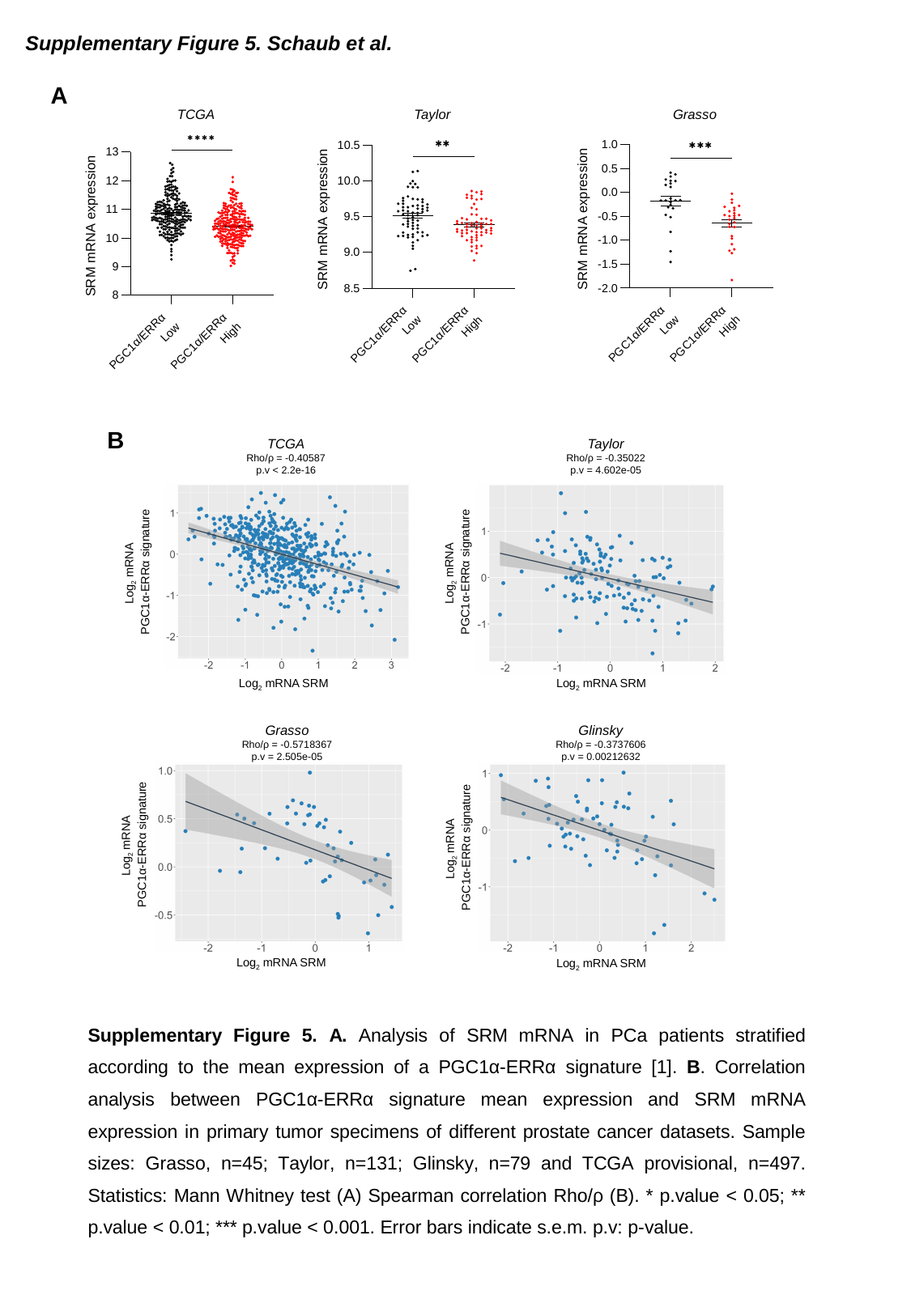

Supplementary Figure 5. Schaub et al.
A
TCGA
Taylor
Grasso
B
TCGA
Rho/ρ = -0.40587
p.v < 2.2e-16
Taylor
Rho/ρ = -0.35022
p.v = 4.602e-05
Log_2_ mRNA
PGC1α-ERRα signature
Log_2_ mRNA
PGC1α-ERRα signature
Log_2_ mRNA SRM
Log_2_ mRNA SRM
Grasso
Rho/ρ = -0.5718367
p.v = 2.505e-05
Glinsky
Rho/ρ = -0.3737606
p.v = 0.00212632
Log_2_ mRNA
PGC1α-ERRα signature
Log_2_ mRNA
PGC1α-ERRα signature
Log_2_ mRNA SRM
Log_2_ mRNA SRM
